# A novel bifunctional peptide VAMP mined from hemp seed protein hydrolysates improves glucose homeostasis by inhibiting intestinal DPP-IV and increasing the abundance of *Akkermansia muciniphila*

**DOI:** 10.1101/2024.07.22.604525

**Authors:** Haihong Chen, Wei Li, Wei Hu, Bing Xu, Yi Wang, Junyu Liu, Chong Zhang, Canyang Zhang, Xizhen Zhang, Qixing Nie, Xinhui Xing

**Affiliations:** Institute of Biopharmaceutical and Health Engineering, Tsinghua Shenzhen International Graduate School, Shenzhen 518055, China; Institute of Biomedical Health Technology and Engineering, Shenzhen Bay Laboratory, Shenzhen 440300, China; Key Laboratory for Industrial Biocatalysis, Ministry of Education, Institute of Biochemical Engineering, Department of Chemical Engineering, Beijing 100084, China; Center for Synthetic and Systems Biology, Tsinghua University, Beijing 100084, China; College of life Science and Technology, Guangxi University, Guangxi, 530004, China; State Key Laboratory of Food Science and Resources, China-Canada Joint Lab of Food Science and Technology, Key Laboratory of Bioactive Polysaccharides of Jiangxi Province, Nanchang University, Nanchang, China

**Keywords:** Hemp seed peptide, VAMP, DPP-IV, hyperglycaemia, *Akkermansia muciniphila*

## Abstract

Discovery of new dipeptidyl peptidase IV (DPP-IV) inhibitory peptides from natural protein resources capable of regulating glucose metabolism in type 2 diabetic populations has been a significant challenge. In this study, we constructed a molecular docking- and machine learning-aided DPP-IV inhibitory peptide library and combined a functional screening approach based on intestinal organoids to discover efficient and new DPP-IV-inhibiting peptides from hemp seed protein hydrolysates. A novel tetrapeptide, VAMP, was then identified to strongly inhibit DPP-IV (IC_50_=1.00 μM *in vitro*), which competitively binds to DPP-IV and improves glucose metabolism *in vivo* with high safety by increasing active glucagon-like peptide-1 (GLP-1) levels in obese mouse models. Interestingly, VAMP specifically promoted the growth and abundance of intestinal *Akkermansia muciniphila in vivo*, at the same time, which was responsible for the improved intestinal barrier function and insulin resistance. Our study demonstrated that the novel bifunctional VAMP can effectively target the DPP-IV-GLP-1 axis and simultaneously regulate the abundance of the gut microbial *A. muciniphila*, to regulate glucose homeostasis, providing a promising nutraceutical and therapeutic tetrapeptide for hyperglycaemia treatment by targeting the gut-microbiata axis.

**Teaser:** VAMP improves glucose metabolism by increasing the active GLP-1 level and promoting the growth of *A. muciniphila* to improve intestinal barrier function.

## Introduction

The disturbance of glucose homeostasis has become a substantial medical problem due to its increasing global prevalence and because disordered glucose metabolism is closely linked with diabetes, obesity, liver disease and several cardiovascular diseases (*1, 2*). Accordingly, exploration of effective approaches to alleviate glucose metabolic disorders has therefore gained increasing attention (*3, 4*). Incretin hormone glucagon-like peptide-1 (GLP-1) has been identified plays an important role in glucose homeostasis via stimulating insulin secretion, improving beta-cell mass, abating gastric emptying, and reducing intestinal peristalsis (*5–7*). However, GLP-1 were rapidly degraded by the enzyme dipeptidyl peptidase IV (DPP-IV), which is a metabolic enzyme with 766 amino acids that can lead to the deactivation of GLP-1 by preferentially cleaving the amino acid residues Pro/Ala and Gly/Ser/Thr at position 2 (*8*). Hence, GLP-1 receptor agonists (GLP1RAs, e.g., liraglutide and exenatide) is popular in the past decades, which mimic the structure of GLP-1 *in vivo* and thus can bind GLP-1R but are not subject to GLP-1 degradation by DPP-IV, hence increasing endogenous GLP-1 concentrations in humans (*9*). However, gastrointestinal adverse effects are widely found in patients treated with GLP1RAs (*10, 11*), even increasing the risk of nutritional deficiencies, acute pancreatitis, pancreatic carcinoma, medullary thyroid carcinoma, diabetic ketoacidosis and diabetic retinopathy (*12*), which further increases the demand for effective, safe and acceptable therapeutic alternative options.

DPP-IV inhibitors are oral antidiabetic drugs that differ from GLP1RAs in terms of their structure and mechanism of action. DPP-IV inhibitors developed so far are small molecules capable of functioning by blocking incretin degradation caused by DPP-IV, hence inhibiting the degradation of GLP-1 and prolonging its half-life (*9*). Numerous DPP-IV inhibitors, including sitagliptin, saxagliptin, linagliptin, alogliptin, have been received market clearance in the past decade. Complete preclinical and clinical evidence supports their roles in the treatment of type 2 diabetes via these chemically synthesized DPP-IV inhibitors. However, limited human data are available to resolve important safety concerns (*13*). Currently, one of the important strategies receiving increasing attention is to discover food-derived bioactive peptides (such as animal- and plant-based protein sources) from natural food protein sources for developing DPP-IV inhibitors to treat diabetes, as food-derived peptides possess various beneficial bioactivities, are generally well metabolized and easily absorbed, and have safer profiles than synthetic drugs (*14, 15*). There have been many reports that food protein-derived peptides contain bioactive sequences capable of effectively inhibiting DPP-IV, as they are diverse in amino acid compositions and sequences (*16*). Currently, approximately 500 peptides with DPP-IV inhibitory IC_50_ values have been retrieved from published papers, but the real-world application of these peptides to functional foods or in basic pharmaceutical peptide development based on effective DPP-IV inhibition has scarcely been reported. In addition, the lack of genomic and proteomic information for many natural protein-containing bioresources hinders the efficient mining of new DPP-IV inhibitory peptides. Currently, the main approaches for the mining of DPP-IV inhibitory peptides are labour-, time- and cost-intensive, including the complicated procedure such as separation, purification, identification and synthesis of target peptides (*17*). Therefore, it is an important challenge to develop a high-throughput integrated mining method which enables effective DPP-IV inhibitory biopeptides to be discovered from natural protein-rich bioresources with an emphasis on systematically regulating glucose homeostasis with high safety that can be used in functional food development and peptide medicine innovation.

Extensive studies have reported the important role of the gut microbiome in regulating host glucose metabolism homeostasis, fat accumulation and mucosal barrier integrity (*10, 18, 19*). Previous studies have shown that obese and diabetic humans and mouse models exhibit disrupted gut microbiota with reduced diversity and impaired mucosal barrier integrity (*20*). Gut *A. muciniphila* was first isolated by Derrien *et al*. from the faeces of a healthy person and has been considered as next probiotics (*21*). The health benefits of *A. muciniphila* on host metabolism have been widely reported, especially on obesity, diabetes, intestinal inflammation and different cancers, mainly based on the regulation of intestinal barrier function (*22–25*). Intervention strategies involving the use of bioactive dietary peptides to modulate the gut microbiota have been proposed to prevent and treat diabetes and related metabolic diseases and are closely associated with the regulation of *A. muciniphila* (*26, 27*). Additionally, another study revealed that treatment with a nonapeptide could alleviate obesity by suppressing appetite and regulating the gut microbiota (*10*). Furthermore, the abundance of *A. muciniphila* was increased by the nonapeptide treatment, which might further inhibit fat absorption by downregulating intestinal Cd36 (*10*). The effects of some dietary proteins or peptides on *A. muciniphila* have been reported, but the discovery and mechanisms of peptides specifically promoting *A. muciniphila* in gut have rarely been reported.

There are numerous challenges associated with synthetic drug molecules, including gastrointestinal instability, high costs, adverse side effects, stringent regulatory compliance, and intense market competition. Hence, the increasing demand for exploring natural bioactive peptides with high safety and efficacy in regulating glucose metabolism via gut-microbita axis necessitates the use of natural protein- and amino acid-rich agricultural resources, as well as the development of an effective bioactive peptide production strategy that is feasible for nutraceutical and pharmaceutical applications. The success of these efforts will largely depend on the rapid identification of bioactive peptide sequences from natural bioresources. Hemp (*Cannabis sativa* L.), a member of the *Cannabaceae* family, originated in Central Asia and is known for its medicinal value and textile applications (*28*). In China, hemp is widely farmed and has long been approved for medicinal use based on the concepts of medicine and food homology. Hemp seeds are rich in proteins (20-25%), which contain various essential amino acids in a desirable ratio (*29*). Hemp seed protein (HSP) hydrolysates exhibit a variety of bioactivities(*3, 29*), our previous study predicted the potential biological activities of HSP-derived peptides by using the BIOPEP-UWM web server and revealed that HSP-derived peptides are rich in DPP-IV inhibitory peptides (*3*). However, the composition of peptides in HSP hydrolysates and their mechanism of action on DPP-IV and glucose reduction have not yet been explored, especially the efficient mining methods of DPP-IV-inhibiting peptides from HSP-derived peptide libraries and the discovery of key target peptides for glucose metabolism regulation.

In this study, we employed molecular docking-, two-layer neural network-based machine learning- and intestinal organoid-based functional screening methods to explore bioactive peptides from HSP capable of inhibiting DPP-IV to regulate glucose homeostasis. Oral administration of a newly mined HSP-derived peptide, VAMP, which has a high content of HSP and biofunctionality, significantly improved the glucoregulatory capacity via the inhibition of intestinal DPP-IV and promotion of *A. muciniphila* growth in obese mice, thereby effectively alleviating hyperglycaemia. Our study demonstrated the feasibility of the integrated mining method of functional peptides from natural bioresources and revealed a new and promising biofunctional peptide with high safety that can be taken orally for the prevention and treatment of hyperglycaemia.

## Results

### Functional screening of DPP-IV-inhibiting peptides from hemp seed protein (HSP)

HSP have recently drawn considerable attention in both scientific and industrial fields because of their excellent nutraceutical value, superior digestibility, low allergenicity, and diverse techno-functional properties. Enzymatic HSP hydrolysates using different enzymes, combined with *in vitro* and *in vivo* functional validation, were utilized to screen the functional enzymatic products derived from the HSP (**Figure 1A**). According to our measurements, the hemp seeds used in this study contained approximately 26% protein (**Figure 1B**), and the HSP consisted of various essential amino acids with a balanced ratio, especially arginine, aspartic acid, glutamic acid, valine, and histidine (**Table S1**). Furthermore, 1184 proteins were identified from the HSP via proteomic analysis. To identify the potential biological functions of the peptides derived from the HSP, the sequences of the 1184 proteins were imported into the BiOPEP-UWM server (https://biochemia.uwm.edu.pl/biopep-uwm/) for functional annotation. Interestingly, we found that HSP-derived peptides might favourably inhibit DPP-IV (44% of the total annotated HSP-derived peptides) and have antioxidative and ACE inhibitory effects (32% and 19%, respectively). These results implied that HSP-derived peptides might have diverse DPP-IV inhibitory effects that could contribute to improving glucose metabolism (**Figure 1C**).

**Figure 1.**
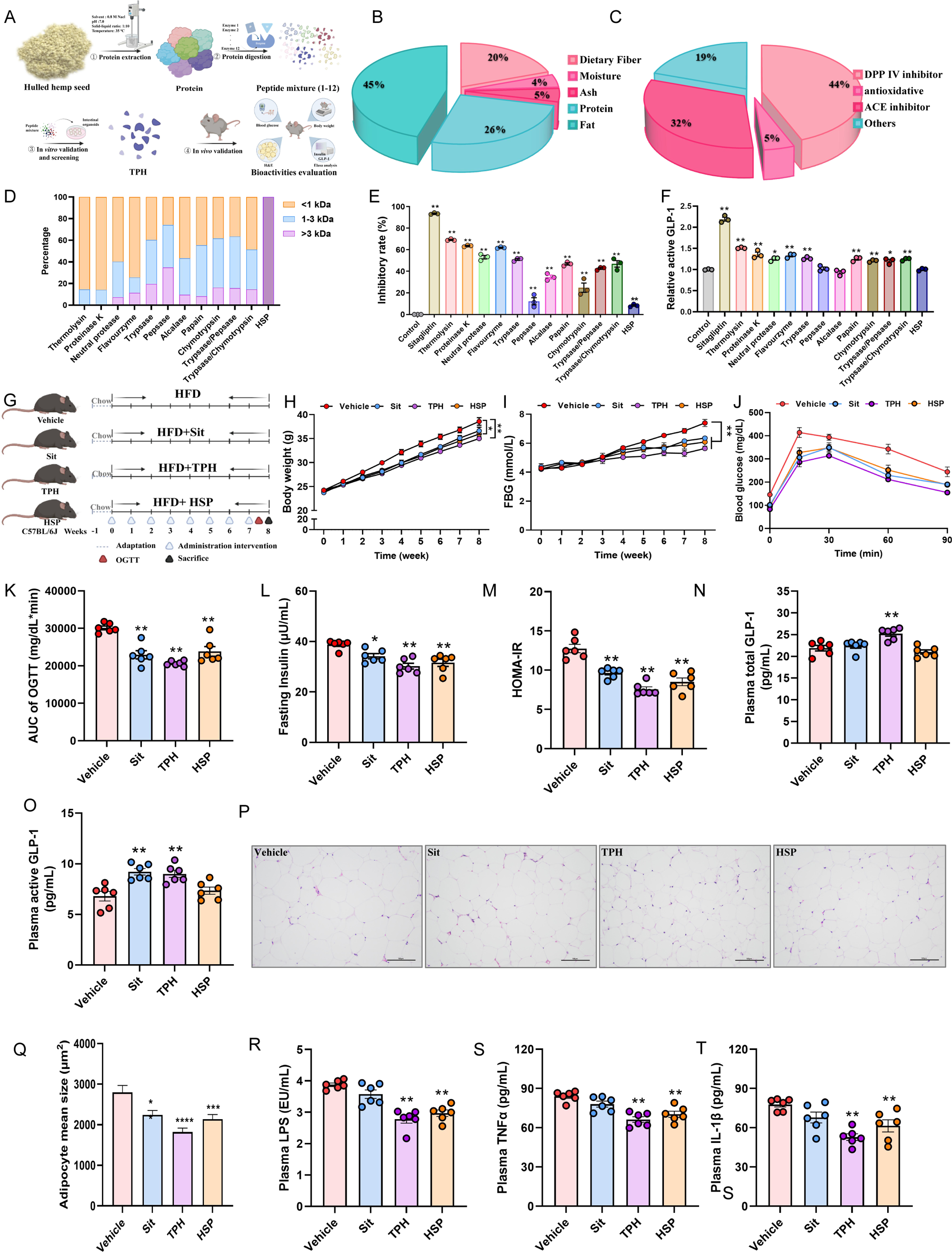
The enzymatic products of HSPs processed by thermolysin inhibit DPP-IV activity and improve glucose metabolism. (A) Strategy for screening enzymes to be used for the enzymolysis of HSPs to obtain hydrolysates that inhibited DPP-IV. Enzymes, including thermolysin, proteinase K, neutral protease, flavourzyme, trypsase, pepsase, alcalase, papain, chymotrypsin, trypsase/pepsase, and trypsase/chymotrypsin, were used for the hydrolysis of HSPs. Subsequently, the hydrolysates from different enzymes were evaluated based on HPGPC analysis, DPP-IV inhibitory activity, and active GLP-1 levels. Finally, animal studies were conducted to validate the hypoglycaemic effects of the selected hydrolysates. (B) The nutritional composition of the hemp seeds was determined. (C) The percentage of peptide sequence numbers in HSPs with various functions is shown. (D) The molecular weight distribution of the HSP hydrolysates generated using different enzymes. (E) The inhibitory rate of different HSP hydrolysates on DPP-IV. (F) The relative concentrations of active GLP-1 in intestinal organoid treated with different HSP hydrolysates. (G) The experimental scheme for (H to T) is shown, n = 6 mice/group. HFD-fed mice were treated with PBS (vehicle group), sitagliptin (sit group), HSP hydrolysates by thermolysin (TPH group), or HSP (HSP group) for 8 weeks by oral gavage. (H) The body weight change, (I) FBG, (J) OGTT curve, (K) AUC of the OGTT, (L) levels of fasting insulin, (M) HOMA-IR index, (N) levels of plasma total GLP-1, and (O) levels of plasma active GLP-1 were determined. (P) Representative H&E images of sWAT deposits (3 mice per group) are shown. Scale bars, 100 μm. (Q) The mean adipocyte size (3 mice per group), (R) plasma LPS level, (S) plasma TNFα level, and (T) plasma IL-1β level were determined. All the data are presented as the mean ± SEM. \**P* < 0.05 and \*\**P* < 0.01 versus the vehicle group.

To investigate the DPP-IV inhibitory spectrum of the HSP-derived peptides, different proteases (including thermolysin, proteinase K, neutral protease, flavourzyme, trypsase, pepsase, alcalase, papain, chymotrypsin, trypsase/pepsase, and trypsase/chymotrypsin) were applied for the enzymolysis of HSP. Thermolysin, proteinase K, and flavourzyme were found to be highly effective for HSP proteolysis, with peptides with molecular weights lower than 1000 Da accounting for 85.52%, 85.82%, and 74.35% of the enzymatic products, respectively (**Figure 1D**). To evaluate the ability of the HSP-derived peptides to inhibit DPP-IV, a human-derived induced pluripotent stem cell (hiPSC)-induced intestinal organoid model was established. Based on the intestinal organoids cultivated with the peptides, we found that hydrolysates obtained via thermolysin (TPH) possessed the strongest DPP-IV inhibitory activity and increased the levels of active GLP-1 to the greatest extent (**Figure 1E-F, Figure S1A-B**).

Because TPH possesses superior DPP-IV inhibitory activity and increases the level of active GLP-1 in intestinal organoids, we next explored the hypoglycaemic activity of TPH *in vivo*. The HFD-fed mice were fed TPH for 8 weeks to investigate the hypoglycaemic effects (**Figure 1G**). We found that TPH treatment significantly suppressed body weight gain and fasting blood glucose (FBG) levels, improved glucose tolerance, and reduced insulin resistance (**Figure 1H-M**). Specifically, TPH treatment significantly increased the total and active GLP-1 in plasma, indicateing that inhibition of the DPP-IV-dependent pathway could be one of the key factors by which TPH improved the glucose metabolism (**Figure 1N-O**). In addition, smaller adipocytes in white adipose tissue, a lower body fat ratio, and decreased plasma NEFA, TG, LDL-c, ALT, and AST levels were observed in the TPH group (**Figure 1P-Q, Figure S1C-J**). Additionally, treatment with TPH decreased plasma endotoxin levels and plasma TNF-α and IL-1β levels (**Figure 1R-T**).

### Mining and discovery of key functional peptides from TPH contributing to DPP-IV inhibition

Because the above results showed that TPH possessed strong DPP-IV inhibitory activity and improved glucose metabolism, we then explored the key active peptide sequences by combining molecular docking and machine learning-based virtual screening, followed by *in vitro* functional evaluation of DPP-IV inhibitory activity (**Figure 2A**). First, we determined the peptide composition of THP via proteomics analysis, and 421 peptides with different amino acid sequences were identified from TPH. Next, molecular docking was performed by using PyRx software, and the peptides were sorted according to their binding affinity. At the same time, a neural network-based virtual screening was also conducted. First, we acquired 3681 compounds with IC_50_ values (against DPP-IV) from the ChEMBL database. Then, a two-layer neural network model was established for the subsequent virtual screening. Then, 2576 compounds were randomly selected as the training set, and 1105 compounds were set as the test set. Finally, models with mean squared errors (MSEs) and mean absolute errors (MAEs) of 1.18 and 0.81, respectively, were established. Subsequently, we made predictions based on the trained model, and the peptides were sorted according to the predicted DPP-IV inhibitory pIC_50_ values. Based on this virtual screening method, dynamic network Venn diagram analysis was performed for the top 100 peptides (tripeptides, tetrapeptides, pentapeptides and hexapeptides) in TPH based on the intensity, affinity and predicted pIC_50_, and the intersection of the three indices containing 24 peptides, namely, VFTPQ, VADW, VAMP, FNPRG, FNVDSE, FPQS, FQL, FLQ, WNVN, WIAVK, PSSQQ, YTPHW, YSYA, YTGD, YQLM, FDGEL, PQNHA, LNAP, YNLP, YQL, LLY, WDSY, WLE and YNL, were selected for further analysis (**Figure 2B-C, Figure S2A-E**). In addition, based on the function-screening results obtained from iPSC-induced intestinal organoids, we found that the peptides VADW, VAMP, FNPRG, FLQ, WIAVK, YQLM, LLY, YSYA, FPQS and WDSY showed relatively strong DPP-IV inhibitory activities (**Figure S2F**). Specifically, 7 peptides were found to have strong DPP-IV inhibition activity based on their affinity for DPP-IV (KD value), IC_50_ values, and relative SPR response (surface plasmon resonance analysis, **Figure 2D**), which were selected for further analysis. The peptides FPQS, YNLP, VAMP, VADW, YQLM, YSYA and LLY were identified as the potential key bioactive peptides in TPH that contributed to hypoglycaemic effects, among which VAMP is the strongest DPP-IV inhibitory peptides (with IC_50,_ relative SPR response and KD values were 1.00 μM, 10.9 RU and 6.89 μM, respectively) (**Figure 1**, **Figure 2D-F**).

**Figure 2.**
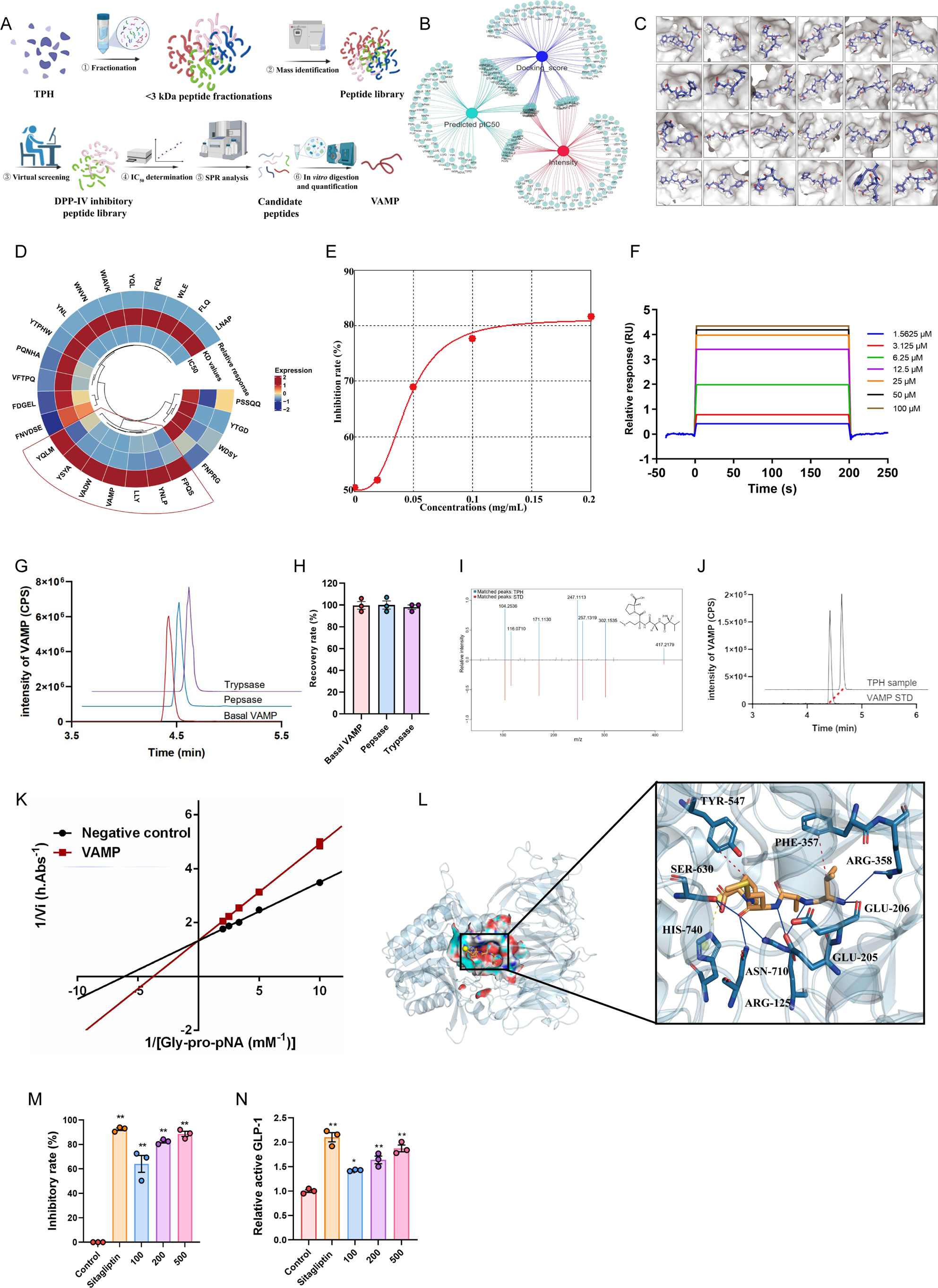
VAMP is an effective peptide inhibitor of DPP-IV. (A) Strategy for the identification of peptides that inhibit DPP-IV. To screen the key active peptides that play vital roles in inhibiting DPP-IV activity, the peptide composition of TPH was analysed using mass spectrometry, and 422 peptides were identified. Next, a virtual DPP-IV inhibitory peptide library was established by using molecular docking and machine learning methods, and 24 peptides were selected for further analysis. To verify DPP-IV inhibitory activity, the IC_50_ values and binding affinities were also evaluated. (B) Dynamic network Venn diagram analysis was performed for the top 100 peptides (tripeptides, tetrapeptides, pentapeptides, and hexapeptides) in TPH based on the intensity, affinity and predicted pIC_50._ (C) The predicted binding model between DPP-IV and the 24 selected bioactive peptides was visualized by PyMOL. (D) Heatmap analysis of the relative response, KD values, and IC_50_ values of the 24 selected peptides was performed. (E) The IC_50_ curve of VAMP. (F) The SPR sensorgrams of click-immobilized DPP-IV toward VAMP at different concentrations are shown. (G) The chromatograms of VAMP after pepsase and trypsase digestion are shown. (H) The recovery rate of VAMP after pepsase and trypsase digestion was determined. (I) The LC–MS/MS data of VAMP from synthetic standards and TPH samples were compared. (J) Extracted ion chromatograms of VAMP in the TPH sample are shown. (K) Lineweaver ‒ Burk plots of DPP-IV inhibition by VAMP are shown. (L) Active site superimposition between VAMP-bound DPP-IV and key residues is labelled with a three-letter code. VAMP (claybank) and residues (navy blue) are shown as sticks, and DPP-IV is shown as a cartoon. Hydrogen bonds are coloured blue, hydrophobic interactions are coloured grey, and salt bridges are coloured yellow. (M) The inhibitory effect of VAMP on DPP-IV was determined. (N) The relative concentrations of active GLP-1 after VAMP treatment in intestinal organoids were measured.

Next, we designed experiments to assess the digestive stability of the seven identified peptides. The results showed that only VAMP, FPQS, and YNLP were readily resistant to the simulated gastric pepsase and trypsase digestion (**Figure 2G-H for VAMP**, **Figure S2G for the other 6 peptides**). We then quantified the levels of these peptides in TPH samples. We detected YSYA, YNLP, VAMP, VADW, and LLY in TPH, among which VAMP was the most abundant (4.58 ± 0.03 μg/mg, **Figure 2I-J, Figure S2H**). To investigate the mechanism by which VAMP inhibits DPP-IV activity, we used Lineweaver–Burk double reciprocal representation methods to analyse the binding mode and showed that VAMP could competitively inhibit DPP-IV activity (**Figure 2K**). PyMOL was utilized to display the molecular interaction between DPP-IV and VAMP, and the results showed that VAMP could form hydrogen bonds with the DPP-IV residues at Arg125, Glu205, Glu206, Arg358, Ser630, and Asn710; establish hydrophobic interactions with residues Phe357 and Tyr547; and establish salt bridges with His740 (**Figure 2L**). A detailed analysis of the interactions between DPP-IV and the other six peptides is shown in **Figure 2SI**. Furthermore, we also found that VAMP inhibited DPP-IV and increased the level of active GLP-1 in a dose-dependent manner in the hiPSC-induced intestinal organoids (**Figure 2M-N**).

### Regulation of glucose metabolism by VAMP via intestinal DPP-IV inhibition

Among the above HSP-derived peptides generated by enzymatic hydrolysis with thermolysin, we found that VAMP exhibited the strongest inhibitory activity against DPP-IV (IC_50_ = 1.00 μM) and was resistant to simulated gastric digestion (**Figure 2G-H**). We firstly tested the safety of VAMP in mice, treatment with VAMP at different dosage (10 mg/kg and 100 mg/kg) did not induce any obvious side effects in the mice (**Figure S3 A-F**). We next treated HFD-fed mice with VAMP for one week to investigate the effects of VAMP on glucose homeostasis through the inhibition of DPP-IV activity (**Figure 3A**). We found that there were no significant differences in body weight or energy intake between the vehicle and VAMP groups (**Figure 3B and C**). However, VAMP treatment significantly decreased intestinal DPP-IV activity (**Figure 3D**) and increased the levels of intestinal and plasma active GLP-1 without altering the intestinal and plasma total GLP-1 levels (**Figure 3E-H**), which was associated with increased levels of glucose-stimulated insulin and a better OGTT (**Figure 3I-K**).

**Figure 3.**
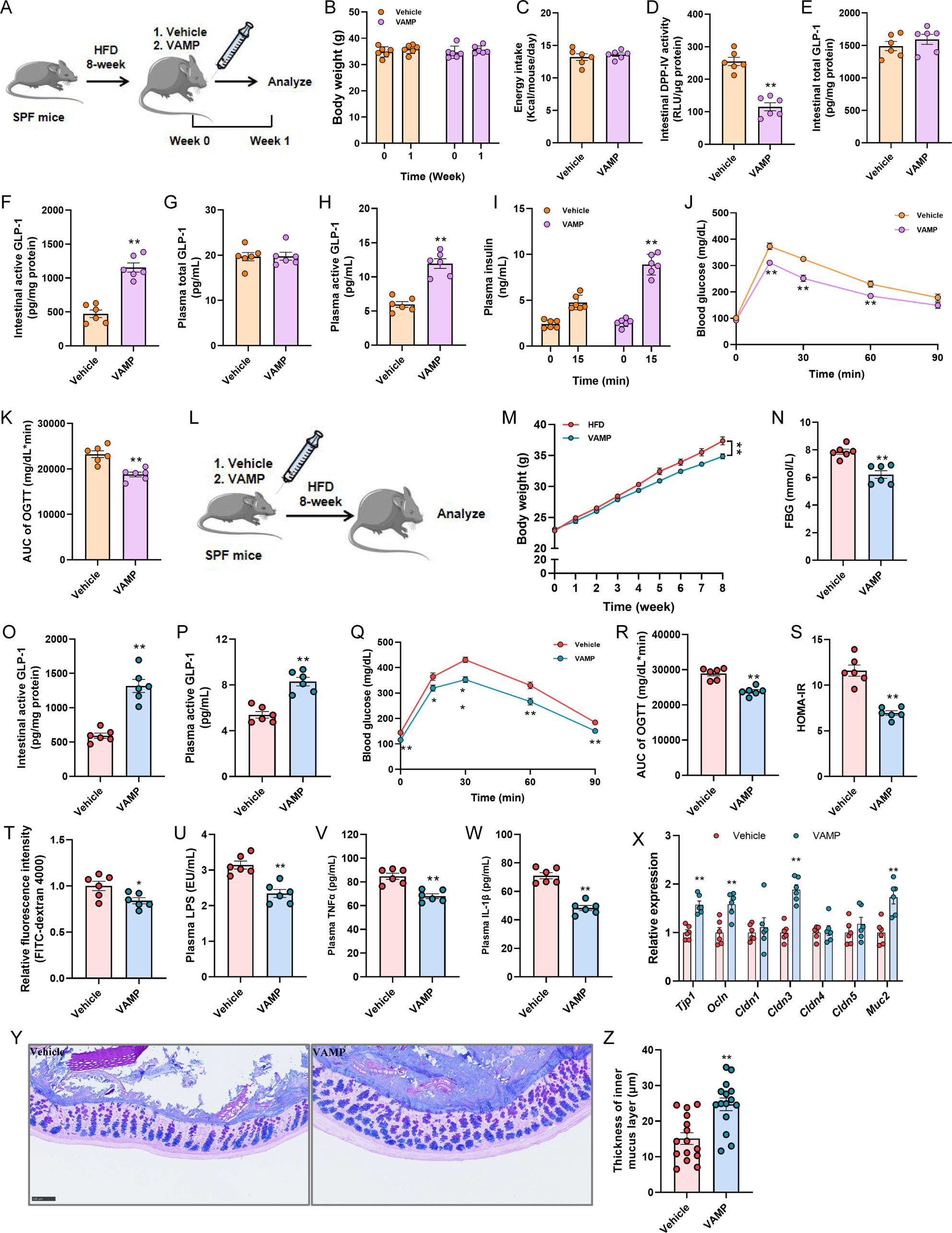
VAMP inhibits intestinal DPP-IV and improves host glucose metabolism. (A) The experimental scheme for (B to K) is shown, n = 6 mice/group. Mice were fed a HFD for 8 weeks, divided into two groups, and treated with PBS (vehicle group) or VAMP (50 mg/kg, VAMP group) for 1 week. (B) The body weights, (C) energy intake, (D) DPP-IV activity in intestinal tissue, (E) intestinal total GLP-1 levels, (F) intestinal active GLP-1 levels, (G) plasma total GLP-1 levels, (H) plasma active GLP-1 levels, (I) glucose-stimulated insulin levels, (J) OGTT curve, and (K) AUC of the OGTT were determined. (L) The experimental scheme for (M to Z) is shown, n = 6 mice/group. HFD-fed mice were treated with PBS (vehicle group) or VAMP (50 mg/kg, VAMP group) for 8 weeks by oral gavage. (M) The body weight change, (N) FBG level, (O) intestinal active GLP-1 levels, (P) plasma active GLP-1 levels, (Q) OGTT curve, (R) AUC of the OGTT, and (S) HOMA-IR index were evaluated. (T) Intestinal permeability was measured by plasma fluorescence intensity after gavage with FITC-dextran 4000. (U) The plasma levels of LPS, (V) TNFα, and (W) IL-1β were determined. (X) The relative expression of *Tjp1*, *Ocln*, *Cldn1*, *Cldn3*, *Cldn4,* and *Cldn5* mRNAs in colonic tissue was determined. (Y) Carnoy-fixed colonic tissue sections were stained with Alcian blue/periodic acid-Schiff. Scale bars, 100 µm. (Z) Blinded colonic mucus layer measurements were made from Alcian blue-stained sections. All the data are presented as the mean ± SEM. \**P* < 0.05 and \*\**P* < 0.01 versus the vehicle group.

We next assessed the effects of VAMP on model mice during the induction of abnormal glucose metabolism. The mice were orally fed a HFD and VAMP for 8 weeks simultaneously (**Figure 3L**). Administration of VAMP significantly decreased mouse body weight, body weight gain, FBG, fat weight/body weight ratio, and the size of adipocytes in white adipose tissue (**Figure 3M-N**, **Figure S3G-J**). As anticipated, supplementation of HFD-fed mice with VAMP for 8 weeks significantly increased the levels of intestinal and plasma active GLP-1 without affecting intestinal or plasma total GLP-1 levels and improved glucose tolerance (**Figure 3O-R**, **Figure S3K-L**); these results were in agreement with the results showing that VAMP could be an effective inhibitor of host DPP-IV. However, we also found that VAMP decreased fasting insulin levels and improved insulin resistance in mice fed a HFD (**Figure 3S**, **Figure S3M**). As the regulation of intestinal barrier function is important for ameliorating insulin resistance and related metabolic diseases, we also examined intestinal barrier function-related parameters. Interestingly, VAMP treatment significantly reduced intestinal permeability and plasma endotoxin levels (**Figure 3T and U**), as well as plasma TNF-α and IL-1β levels (**Figure 3V and W**), compared with those in HFD-fed control mice. In addition, VAMP was found to upregulate the expression of intestinal *Tjp1*, *Ocln*, *Cldn3*, and *Muc2*, along with the increased thickness of mucus layer (**Figure 3X-Z**). We also assessed the effects of VAMP in *ob/ob* mice as a genetic model of obesity (**Figure S3N**). There was no significant difference in body weight or energy intake, but the FBG steadily decreased with VAMP treatment (**Figure S3O-Q**). VAMP also suppressed intestinal DPP-IV activity and increased the levels of intestinal and plasma active GLP-1, which were associated with the increased levels of glucose-stimulated insulin and improved glucose tolerance (**Figure S3R-Y**). Collectively, these results indicated that VAMP could ameliorate glucose intolerance via the inhibition of intestinal DPP-IV and improve intestinal barrier function in obese mice.

### Biofunctional effects of VAMP on the gut microbiota and specific enhancement of *A. muciniphila* abundance

Dysbiosis of the gut microbiota is associated with the onset and progression of insulin resistance, and we found that VAMP could alleviate insulin resistance and the inflammatory response. Consequently, in addition to inhibiting intestinal DPP-IV, we also hypothesized that VAMP could affect the composition and function of the gut microbiota. We first determined the site of VAMP activity by metabolic kinetics. According to the metabolic kinetics results, VAMP showed a bioavailability of 10.54% after *i.v.* and *p.o.* once in mice, indicating that VAMP has poor oral absorption (**Figure 4A**). Furthermore, 74.18% of VAMP remained in the intestinal content after gavage at the time of peak plasma concentration (**Figure 4B**), implied that in addition to intestinal DPP-IV inhibition, VAMP might interact with the gut microbiota to exert beneficial effects on the intestinal lumen.

**Figure 4.**
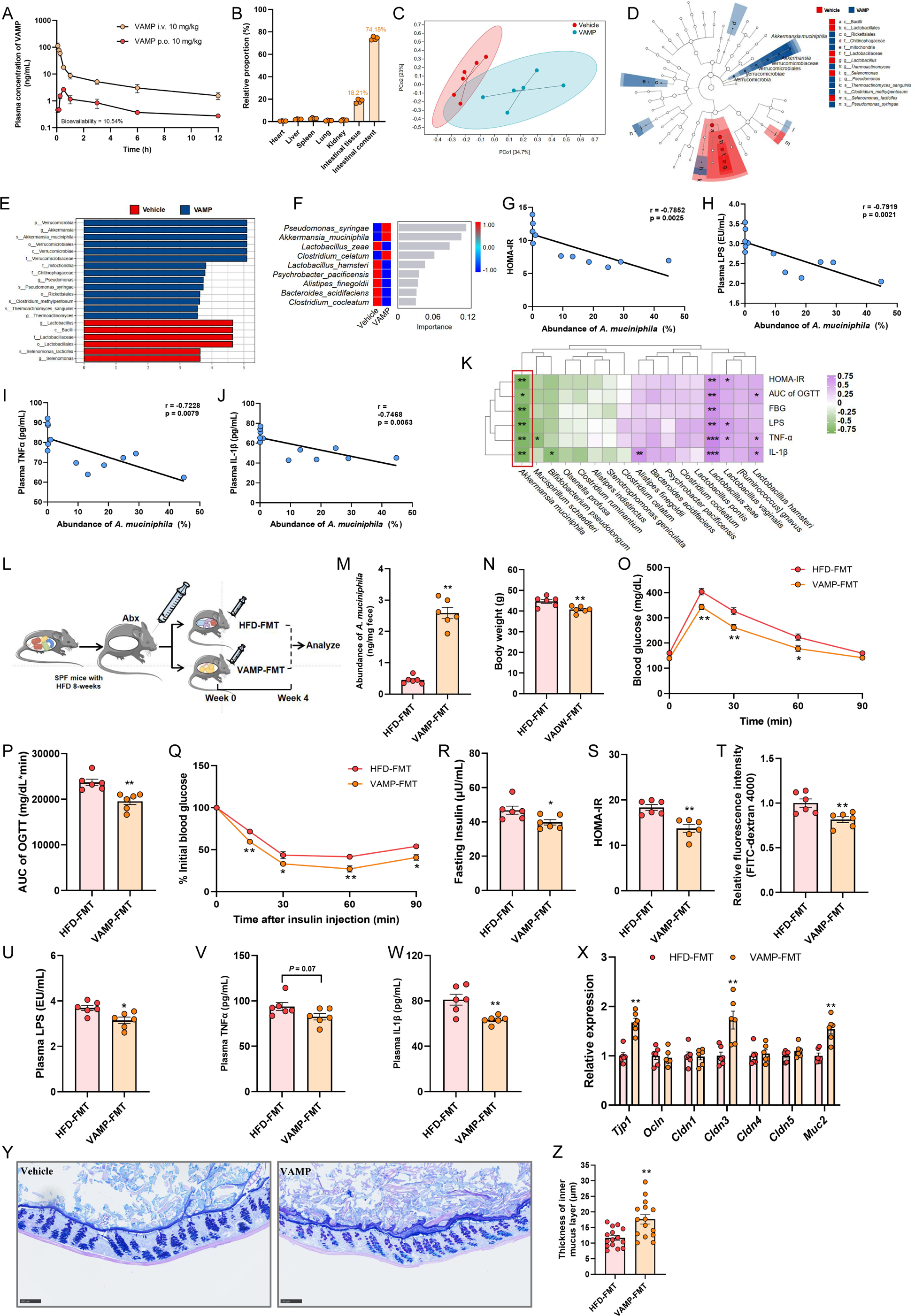
VAMP regulates the gut microbiota and improves host glucose metabolism. (A) The blood pharmacokinetics of 10 mg/kg VAMP administered by injection (*i.v.*) or gavage (*p.o.*) over time were evaluated. n = 4 mice/group. (B) The tissue distribution of VAMP at the time of peak plasma concentration (30 min) after the administration of 10 mg/kg VAMP by gavage was determined; n = 4 mice/group. (C-K, related to Figure 3L) HFD-fed mice were treated with PBS (vehicle group) or VAMP (50 mg/kg, VAMP group) for 8 weeks by oral gavage. n = 6 mice/group. (C) Principal coordinate analysis (PCoA) was performed using the Bray‒Curtis distance. (D-E) Taxonomic cladograms were generated by LEfSe analysis. The blue colour indicates enriched taxa in the VAMP group, and the red colour indicates enriched taxa in the vehicle group. The size of each circle is proportional to the taxon’s abundance. (F) The change in bacterial abundance at the species level after VAMP treatment was determined via random forest analysis. (G-J) Correlations of *A. muciniphila* abundance with the HOMA-IR score, plasma LPS level, plasma TNFα level, and plasma IL-1β level were performed. Correlations between variables were assessed by linear regression analysis. The linear correction index R and P values were calculated. (K) Spearman correlation analysis between the gut microbiota and inflammation- and glucose metabolism-related parameters was performed. (L) The experimental scheme for M to Z is shown, n = 6 mice/group. Mice were fed a HFD for 8 weeks and treated with Abx for 1 week. Then, faecal homogenates from VAMP-treated or untreated HFD-fed mice were orally transferred to Abx-treated recipient mice. (M) *A. muciniphila* abundance in the faeces was assessed by qPCR. (N) The body weights, (O) OGTT curve, (P) AUC of the OGTT, (Q) ITT, and (R-S) fasting insulin levels and HOMA-IR index of HFD-FMT and VAMP-FMT mice were evaluated. (T) Intestinal permeability was measured by plasma fluorescence intensity after gavage with FITC-dextran 4000. (U) The plasma levels of LPS, (V) TNFα, and (W) IL-1β were determined. (X) The relative expression of *Tjp1*, *Ocln*, *Cldn1*, *Cldn3*, *Cldn4,* and *Cldn5* mRNAs in colonic tissue was evaluated. (Y) Carnoy-fixed colonic tissue sections were stained with Alcian blue/periodic acid-Schiff. Scale bars, 100 µm. (Z) Blinded colonic mucus layer measurements of Alcian blue-stained sections were performed. All the data are presented as the mean ± SEM. \**P* < 0.05 and \*\**P* < 0.01 versus the HFD-FMT group.

To investigate the influence of gut microbiota composition induced by VAMP treatment, colonic microbiota was analysed by 16S rRNA gene sequencing. There was no significant differences in α diversity (observed species, Chao1, and Shannon indices) between HFD and VAMP group (**Figure S4A**). However, treatment with VAMP significantly influenced the composition of colonic microbiota based on the analysis of β diversity (**Figure 4C, Figure S4B and C**). At the phylum level, VAMP increased the abundance of Verrucomicrobia and decreased the abundance of Actinobacteria (**Figure S4D**). At the genus level, we found that the abundance of *Akkermansia* was significantly increased by VAMP treatment, along with a decreased abundance of *Lactobacillus* and *Streptococcus* (**Figure S4E and F**). Specifically, LEfSe method revealed that the VAMP group was characterized by *A. muciniphila*, *Pseudomonas syringae*, *Clostridium methylpentosum*, and *Thermoactinomyces sanguinis*, among which *A. muciniphila* had the highest abundance (**Figure 4D and E**, **Figure S4G**), and the results of random forest classification also highlighted that the increased abundance of *A. muciniphila* by VAMP was significant for the classification of the two groups (**Figure 4F**). More importantly, the abundance of *A. muciniphila* was negatively correlated with HOMA-IR, plasma endotoxin levels, TNFα levels, and IL-1β levels (**Figure 4G-K**). With the peptides in TPH characterized by a relatively high abundance of VAMP, we also investigated the effect of TPH on the composition of the gut microbiota. TPH did not influence α-diversity but did change the composition of the gut microbiota in HFD-fed mice (**Figure S4H and I**). As anticipated, TPH also significantly increased the abundance of *A. muciniphila* in the intestine, suggesting that VAMP might be a major effector in promoting the growth of *A. muciniphila* (**Figure S4J-L**). In contrast, although sitagliptin effectively improved glucose metabolism in obese mice, the hypoglycaemic agent had no effect on the regulation of *A. muciniphila* (**Figure S4M-P**).

To determine whether the VAMP-regulated gut microbiota (especially the increase in abundance of *A. muciniphila*) also mediated the metabolic protective effects of VAMP, we performed FMT experiments to determine whether the phenotype resulting from VAMP treatment could be transferred (**Figure 4L**). The FMT efficiency was verified by detecting the enrichment of *A. muciniphila* in mice that received FMT from VAMP-treated mice (**Figure 4M**). There was no significant difference in energy intake between the two groups (**Figure S4Q**), whereas VAMP-FMT mice had significantly lower body weights than HFD-FMT mice (**Figure 4N**). VAMP-FMT mice exhibited improved glucose tolerance and insulin sensitivity (**Figure 4O-S**). Consistent with the results for VAMP-treated HFD mice, VAMP-FMT-fed mice had reduced intestinal permeability, and plasma endotoxin, TNF-α and IL-1β levels (**Figure 4T-W**). Following FMT, intestinal *Tjp1*, *Cldn3*, and *Muc2* were upregulated in VAMP-FMT group, and the thickness of the mucus layer increased (**Figure 4X-Z**). Moreover, the plasma TG level was also decreased in the VAMP-FMT group, with no effects on the plasma TC or NEFA levels (**Figure S4R-T**).

### VAMP enabling intestinal barrier function improvement to exert hypoglycaemic effects via promoting growth of *A. muciniphila*

As we found that VAMP increased the abundance of *A. muciniphila in vivo*, we wondered whether *A. muciniphila* could also be a crucial factor capable of mediating the metabolic protective effects of VAMP. We first investigated the influence of VAMP on the gut microbiota. In the faeces-derived *in vitro* microbial communities, VAMP was degraded by the gut microbiota (**Figure 5A**). There was no significant difference between the VAMP-treated and vehicle-treated groups in terms of α-diversity, but VAMP influenced β-diversity after *in vitro* fermentation (**Figure S5A and B**). The LEfSe results identified *Akkermansia* as a distinguishable genus whose abundance increased after VAMP treatment, which was confirmed by heatmap analysis (**Figure 5B and C**). Furthermore, we detected VAMP after oral gavage, suggesting that VAMP could influence the composition and abundance of the gut microbiota (**Figure 5D**). As VAMP can promote the growth of *A. muciniphila* both *in vitro* and *in vivo*, we next investigated whether *A. muciniphila* could improve glucose metabolism and insulin resistance. Consistent with previous findings, gavage with *A. muciniphila* decreased body weight and improved glucose tolerance, insulin sensitivity, and intestinal barrier function (**Figure S5C-L**).

**Figure 5.**
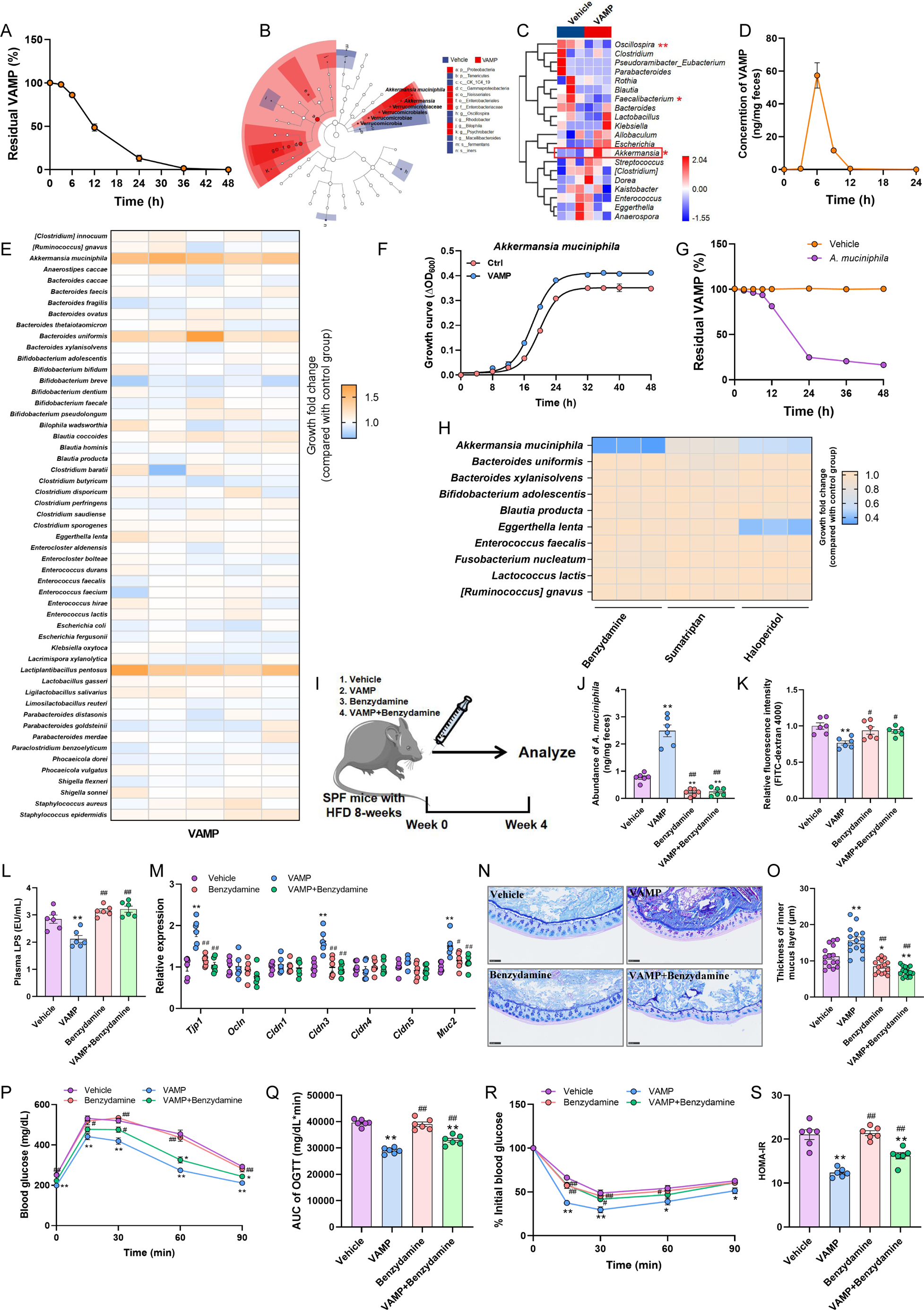
VAMP improves host glucose metabolism by promoting the expansion of *A. muciniphila*. (A-C) Mouse faeces-derived *ex vivo* communities were cultured in BHI media supplemented with VAMP (200 μM) or PBS control and then grown anaerobically for 48 h. (A) The level of remaining VAMP in the medium during incubation with faecal bacteria for 48 h (n = 3 independent incubations) was determined. (B) Taxonomic cladograms were generated by LEfSe analysis. Red indicates enriched taxa in the VAMP group, and blue indicates enriched taxa in the vehicle group. The size of each circle is proportional to the taxon’s abundance. (C) A heatmap was used to compare the gut microbiota between the vehicle and VAMP groups at the genus level. (D) Mice were gavaged with VAMP (20 mg/kg), and the faeces were collected to measure the level of faecal VAMP. (E) Heatmap showing the effect of VAMP (50 μM) incubation on the growth of gut bacteria at the single-strain culture level. VAMP or control PBS was added to the medium and incubated with the indicated strains at the logarithmic phase (inoculum size of 5%), after which the strains were grown anaerobically for 48 h. The values represent the ratio of the OD_600_ between the VAMP group and the control group, n = 5 biological replicates. (F) Growth curve of *A. muciniphila* cultured in BHI media supplemented with VAMP (50 μM) or PBS (control) and then grown anaerobically for 48 h; n = 3 biological replicates. (G) The level of remaining VAMP following incubation with VAMP and *A. muciniphila* for 48 h (n = 3 independent incubations). (H) Heatmap showing the effect of different drugs on the growth of *A. muciniphila*. Drugs at 50 μM or PBS were added to the BHI media and incubated with *A. muciniphila* or other strains at the logarithmic phase (inoculum size at 5%), and the strains were then grown anaerobically for 48 h. Each value represents the ratio of the OD_600_ value between the treatment group and the control group, n = 3 biological replicates. (I) The experimental scheme for J to S is shown, n = 6 mice/group. Mice were fed a HFD for 8 weeks and then treated with PBS (vehicle group), VAMP (50 mg/kg, VAMP group), benzydamine (50 mg/kg, benzydamine group), or VAMP plus benzydamine (50 mg/kg, VAMP+benzydamine group) three times per week for 4 weeks. (J) *A. muciniphila* abundance in faeces was assessed by qPCR. (K) Intestinal permeability was measured by determining the plasma fluorescence intensity after FITC-dextran 4000 gavage. (L) The plasma LPS level was determined. (M) The relative expression of *Tjp1*, *Ocln*, *Cldn1*, *Cldn3*, *Cldn4,* and *Cldn5* mRNAs in colonic tissue was determined. (N) Carnoy-fixed colonic tissue sections were stained with Alcian blue/periodic acid-Schiff. Scale bars, 100 µm. (O) Blinded colonic mucus layer measurements from Alcian blue-stained sections were performed. (P) The OGTT curve, (Q) AUC of the OGTT, (R) ITT curve, and (S) HOMA-IR index of different groups were evaluated. All the data are presented as the mean ± SEM. \**P* < 0.05 and \*\**P* < 0.01 versus the vehicle group; #*P* < 0.05 and ##*P* < 0.01 versus the VAMP group.

After observing the changes in the gut microbiota *in vivo* and *in vitro*, we further explored the effects of VAMP at the level of single bacterial strains. By incubating VAMP with different bacterial strains belonging to 54 different species, we found that VAMP significantly increased the growth of *A. muciniphila*, *Bacteroides uniformis*, and *Lactiplantibacillus pentosus* (**Figure 5E**). According to the bacterial growth curves in liquid media, VAMP significantly promoted the growth of *A. muciniphila* (**Figure 5F**). We also found that *A. muciniphila* could degrade VAMP, suggesting that VAMP could serve as a substrate or inducer for *A. muciniphila* growth (**Figure 5G**). Specifically, we found that VAMP also promoted the growth of *Bacteroides uniformis* but failed to stimulate the growth of other strains of *Bacteroides* (**Figure 5E**). We validated this phenomenon by bacterial growth curves; these results indicated that the growth-promoting effects of VAMP on bacteria are specific (**Figure S5M-N**).

To investigate the role of *A. muciniphila* in glucose metabolism and insulin resistance during VAMP intervention, we developed an *A. muciniphila* ‘deletion’ method in the present study. A previous research exploring the inhibitory effects of a thousand different drugs on different gut commensal bacteria provided us with a list of drugs that specifically inhibited *A. muciniphila*. By supplementing the medium with these drugs that can selectively inhibit the growth of *A. muciniphila*, we found that benzydamine hydrochloride (abbreviated as benzydamine) could specifically inhibit growth of *A. muciniphila*; with no inhibitory or promotional effect on other bacteria (**Figure 5H**). We then orally administered PBS (vehicle), only VAMP, only benzydamine, or VAMP plus benzydamine to HFD-induced obese mice for the last 4 weeks (**Figure 5I**). VAMP treatment in obese mice significantly increased the abundance of *A. muciniphila,* as determined by an absolute quantitative assay by qPCR, and benzydamine treatment reduced the abundance of *A. muciniphila* to an extremely low level (**Figure 5J**). VAMP treatment significantly decreased the body weight of obese mice, but these effects disappeared after simultaneous benzydamine administration (**Figure S5O**). Consistent with the increased abundance of *A. muciniphila*, supplementation with VAMP improved intestinal barrier function and alleviated endotoxaemia and the inflammatory response (**Figure 5K-O, Figure S5P and Q**). However, the use of the *A. muciniphila* ‘remover’ benzydamine counteracted the beneficial effect of VAMP on intestinal barrier function, metabolic endotoxaemia, and inflammation (**Figure 5K-O, Figure S5P and Q**). In contrast, although VAMP treatment improved the FBG, glucose intolerance, and insulin resistance under *A. muciniphila*–deficient conditions (VAMP + benzydamine group) compared with those in the vehicle group, the presence of *A. muciniphila* further improved glucose metabolism in obese mice (**Figure 5P-S, Figure S5R**). These results indicated that the growth-promoting effects of VAMP on *A. muciniphila* are another important effect of the peptide on glucose metabolism.

## Discussions

Many antidiabetic drugs have been used in the clinic for glycaemic control, but it is still difficult to simultaneously meet all the requirements, including satisfactory blood glucose level management, few side effects, and high patient compliance (*30*). GLP-1 could regulate blood glucose by promoting the glucose-dependent secretion of insulin, but which is rapidly degraded by DPP-IV in the host (*31*). GLP-1 receptor agonists and DPP-IV inhibitors have gained popularity in the pharmaceutical market, and food-derived peptides are reported to have multiple biological functions (*32, 33*). Many peptides enzymatic hydrolysis from dietary proteins have been shown to exhibit favorable DPP-IV inhibitory activity in glycemic control, with fewer adverse effects than synthetic drugs that are used for diabetes treatment. In this study, we found that HSP hydrolysates inhibited DPP-IV, and the inhibitory activity varied with the enzymes utilized (**Figure 1E-F**). Hydrolysates obtained via thermolysin hydrolysis possessed the strongest DPP-IV inhibitory activity and the greatest increase in active GLP-1 levels. Inhibition of DPP-IV by HSP-derived VAMP is associated with an increase in active GLP-1 levels and improved glucose tolerance, which has been demonstrated in different animal models (HFD-induced obese mice and *ob*/*ob* mice). These results indicate that harnessing the DPP-IV-dependent pathway by VAMP is an important mechanism to regulate glucose homeostasis *in vivo*.

The enzymatic hydrolysis of proteins is a commonly employed technique in the mining of bioactive peptides. Whereas the traditional method requires the complicated procedure such as separation, purification, identification and synthesis of target peptides, all of which are resource-intensive in terms of labor, time, and finances (*29*). We identified VAMP as a bioactive tetrapeptide with DPP-IV inhibitory effects from thermolysin hydrolysis based on a series of high-throughput screening methods (including multiomics, molecular docking, and machine learning technologies). Compared with traditional mining methods, high-throughput screening methods possess many advantages, including high mining efficiency, strong targeting ability, and low false-positive rates, and this method can be used for the mining of biopeptides produced from various natural protein sources. DPP-IV inhibitors have no intrinsic glucose-lowering activity, so their efficacy as hypoglycaemic agents is directly related to their ability to inhibit DPP-IV activity and is mediated through the effects of the substrates they protect. Among these factors, the affinity for and inhibition of DPP-IV are the most important (*34*). In addition, gastrointestinal digestive stability should also be considered, as peptides should have sufficient resistance to digestion by proteases that exist in the gastrointestinal tract (pepsase and trypsase are two of the most important) to enable DPP-IV inhibition. We found that the peptide VAMP possessed high gastrointestinal digestive stability, which could be attributed to the absence of cleavage sites for pepsase and trypsase in VAMP, as pepsase preferentially cleaves sites, including Phe-|-Xaa, Tyr-|-Xaa, Leu-|-Xaa and Trp-|-Xaa, and trypsase-specific cleavage sites, including Arg-|-Xaa and Lys-|-Xaa (Xaa is Phe, Trp or Tyr) (*35, 36*). The peptides exert DPP-IV inhibitory activity by binding at the active site and/or outside the catalytic site of DPP-IV, the inhibitory activity of different peptides varies greatly, and the peptides generally show different inhibitory modes, including competitive, uncompetitive, noncompetitive and mixed-type modes (*37*). The peptide VAMP could competitively bind to DPP-IV, which was in line with a report showing that a competitive mode of action is observed for most food-derived peptides that have a proline at their P1 position.

Recently, there has been an increasing amount of evidences on the role of the gut microbiota in various metabolic diseases, and manipulation of the gut microbiota might be a novel therapeutic strategy. Bioactive peptides have been reported to modify the composition of the gut microbiota, including by inhibiting the growth of opportunistic pathogens and promoting that of some beneficial commensal bacteria (*10, 38, 39*). In our study, VAMP was resistant to degradation by pepsase and trypsase. Interestingly, after VAMP treatment, a large amount of VAMP was detected in the intestinal content, and the eight-week VAMP intervention was associated with decreased gut permeability, increased thickness of the mucus layer, and increased gene expression of tight junction proteins in HFD-fed mice, indicating a potential interaction with the gut microbiota. Specifically, our hypothesis was validated with an FMT experiment (**Figure 4L-Z**), which demonstrated that the VAMP-induced alterations in the gut microbiota improved glucose metabolism in obese mice.

*A. muciniphila* plays an important role in the maintaining of intestinal barrier integrity, thereby modulating the host inflammatory response (*22*). The decreased abundance of *A. muciniphila* has been reported in numerous metabolic diseases. In the present study, both *in vitro A. muciniphila* fermentation experiments and *in vivo* animal experiments showed that VAMP promoted the growth of *A. muciniphila*, whereas sitagliptin (a traditional DPP-IV inhibitor) did not alter the abundance of the bacteria. *A. muciniphila* has been shown to enhance intestinal barrier integrity by replenishing mucus layer thickness and upregulating the production of antimicrobial peptides in obesity and related metabolic diseases (*40*). High-fat diet was found to decrease mucus thickness and enhance intestinal permeability, which contributes to the increased circulating LPS and pro-inflammatory factor levels, subsequently promoting insulin resistance through Toll-like receptor-4 (TLR-4) activation in the host (*41–43*). FMT and supplementation with VAMP significantly increased the abundance of *A. muciniphila,* which subsequently improved intestinal barrier function, the inflammatory response and insulin resistance (**Figure 3M-Z**, **Figure 4C-Z**).

The mucin present in the mucus layer consists of a peptide backbone extensively adorned with O-linked glycans (*44*). The glycans are composed of a plethora of sugar groups that contain mannose, galactose, fucose, and N-acetylhexosamines (such as N-acetylglucosamine and N-acetylgalactosamine), among which only the N-acetylhexosamines can be used for the growth of *A. muciniphila* (*45, 46*). Notably, only a few other substrates were found to stimulate the growth of *A. muciniphila* in pure culture (*22*). In our study, we found that VAMP could promote the growth of *A. muciniphila* but had little impact on other commensal bacteria in the gut. Numerous studies was found supplementation with bioactibe dietary compounds such as polysaccharides or polyphenols could increase the abundance of *A. muciniphila*. However, due to the complex interaction between dietary compounds and the gut microbiota, it is not clear whether the effects are directly promotion of cross-feeding. Based on a previous study exploring the inhibitory effects of 1000 drugs on the bacterial community, we used an *A. muciniphila* “remover” that had little impact on the overall bacterial community (*47*). We found that the beneficial effect of VAMP on glucose metabolism was significantly decreased in the presence of benzydamine hydrochloride. These results obviously demonstrated that the increased abundance of *A. muciniphila* plays an important role in glucose metabolism during VAMP treatment.

In summary, in our study, we identified the bifunctional peptide VAMP from HSPs that can inhibit DPP-IV, proved that this tetrapeptide improves glucose metabolism via the inhibition of intestinal DPP-IV in obese mice. At the same time, VAMP also specially promotes the growth of *A. muciniphila* among gut microbiota, leading to the alleviation of insulin resistance by improving intestinal barrier function. Our study reported a new and promising safe dietary peptide that can be taken orally for the prevention and treatment of hyperglycaemia by targeting gut-microbiota axis

## Methods

### Chemicals and reagents

Human DPP-IV was obtained from QiZheng Biotech (Shanghai, China). The DPP-IV-Glo^TM^ Protease Assay Kit (G8351) was purchased from Promega (Madison, WI). An active GLP-1 ELISA kit was purchased from Millipore (Millipore, cat# EGLP-35K). Total GLP-1, LPS, TNFα, IL-1β and NEFA kits were purchased from COIBO BIO Technology (Shanghai, China). An intestinal organoid culture kit was purchased from STEMCELL Technologies (Canada). The peptides VFTPQ, VADW, VAMP, FNPRG, FNVDSE, FPQS, FQL, FLQ, WINVN, WIAVK, PSSQQ, YTPHW, YSYA, YTGD, YQLM, FDGEL, PQNHA, LNAP, YNLP, YQL, LLY, WDSY, WLE and YNL (> 98% purity) were synthesized by Bankpeptide Inc. (Anhui Province, China). Series S sensor chips (CM5), HEPES buffer, an amine coupling kit, sodium acetate, N-hydroxysuccinimide (NHS), ethanolamine-HCl, 50 mM NaOH and 10 mM glycine were obtained from GE Healthcare Life Sciences (Uppsala, Sweden). Sitagliptin was purchased from Selleck (Houston, TX). Krebs-Ringer bicarbonate-HEPES buffer (KRBH) was obtained from Shanghai Yuanye Bio-Technology Co., Ltd. (Shanghai, China).

### Nutritional composition analysis of hemp seeds

Hemp seeds (*Cannabis sativa* L.) were obtained from Bama Miao Autonomous County and the Guangxi Zhuang Autonomous Region, China. The dietary fiber, moisture, ash, protein, fat, and amino acid composition of the hemp seeds were determined according to the Chinese National Standard GB 5009.88-2014, GB 5009.3-2016, GB 5009.4-2016, GB 5009.5-2016, GB 5009.6-2016 and GB 5009.124-2016, respectively.

### Functional annotation of hemp seed peptides

The simulation of the protease cleavage sites of hemp seed proteins and peptide functional annotation were performed according to our previous methods (*3*). In brief, 1184 proteins identified in our previous study were imported into PeptideMass (https://web.ExPASy.org/peptide_mass/), and simulation cleaves were performed with trypsase, chymotrypsin, trypsase/chymoptrypsin, pepsase, proteinase K and thermolysin. The cleaved peptides were selected in the range of 0-3000 Da. Following functional annotation, the peptides were imported into the BIOPEP-UWM server (http://www.uwm.edu.pl/biochemia/index.php/pl/biopep/).

### Preparation of protein hydrolysate

The preparation of protein hydrolysates was performed according to our previous methods (*48*). Briefly, hemp seed oil was removed with ethanol, protein was extracted with 0.8 mol/L NaCl aqueous solution (pH 7.0) following the enzymatic hydrolysis of hemp seed protein, and the details of the enzymatic hydrolysis conditions are displayed in **Table S2**.

### Molecular weight distribution of hemp seed protein hydrolysates

The homogeneity and distribution of the enzymolysis solutions were evaluated using high-performance gel permeation chromatography (HPGPC) according to our previous method (*49*). In brief, an HPLC instrument (Agilent 1260) equipped with a TSK-Gel G2000SWXL (300 mm×7.8 mm) column was used for data acquisition. The mobile phase was acetonitrile:water:trifluoroacetic acid (30:70:0.1), the wavelength was set at 220 nm, the flow rate was 0.5 mL/min, and the column temperature was 30 °C. Cytochrome C, aprotinin, VPSGPLGPEGPR, glutathione, and glutamate were used as standards to plot a standard curve under the described chromatographic conditions.

### Evaluation of DPP-IV inhibition by hemp seed protein hydrolysates

iPSC-induced intestinal organoids were established according to the manufacturer’s instructions (STEMdiff^TM^ Intestinal Organoid kit). After the intestinal organoids had fully established (passaged at 3-10 generations), the organoids were treated with 500 μg/mL of hydrolysates for 24 h, the mixtures were centrifuged at 500 × g for 5 min, and the supernatants were used to determine the inhibition of DPP-IV and the concentrations of active GLP-1 by using the DPP-IV-Glo^TM^ Protease Assay Kit and GLP-1 ELISA Kit, respectively.

### Identification of bioactive peptides in TPH

To identify bioactive peptides in TPH, the <3 kDa fraction of TPH was collected. A nanoLC-MS/MS (Thermo Scientific, Waltham, MA) was used for raw data acquisition. Briefly, peptides were dissolved in mass spectrometry loading buffer and separated on a Waters ACQUITY UPLC® BEH C18 column (2.1 mm × 50 mm, 1.7 μm particle size). The gradient elution consisted of 0.1% formic acid in water (solution A) with formic acid in acetonitrile (solution B) as the mobile phase at a flow rate of 250 nL/min. The gradient program for the binary mobile phase system was as follows: 0-2 min, 3-7% B; 2-52 min, 7-22% B; 52-62 min, 22-35% B; 62-64 min, 35-90% B; and 64-84 min, 90-90% B. The m/z scan range was 200 to 2000. MaxQuant 1.5.3.17 software was used for raw data processing.

### Molecular docking of the peptides with DPP-IV

Molecular docking was performed according to our previously methods (*3*). In briefly. PyRx version 0.9 (Autodock 4.0) software was applied to the potential binding capacity evaluation The 3D crystal structure of 1NU8 complexes was selected as protein receptor. The DPP-IV-peptide interactions were evaluated based on the binding affinity.

### Machine learning-based pIC_50_ prediction

A sample two-layer neural network was built to obtain the predicted pIC_50_ values. Briefly, a subset of the dataset containing 3681 DPP-IV inhibitors was acquired from the ChEMBL database (https://www.ebi.ac.uk/chembl/). Next, we established a model according to the Keras workflow, which included preparing the data, defining the model, compiling the model, fitting the model, and evaluating the model and predictions of the peptides (**Figure S2C**). The mean squared error (MSE) and mean absolute error (MAE) were used for the evaluation of the established model (*50–54*).

### Selection of potential bioactive peptides

To screen the potential bioactive peptides, dynamic network Venn diagram analysis was performed for the top 100 peptides (tripeptides, tetrapeptides, pentapeptides and hexapeptides) in TPH based on the intensity, affinity and predicted pIC_50._

### Evaluation of DPP-IV inhibitory activity for the selected peptides

A DPP-IV-Glo^TM^ Protease Assay Kit was used to determine the inhibitory effect of the peptides on DPP-IV. The experiment was performed according to the instructions. The mode of DPP-IV inhibition was analysed according to our previously methods (*3*).

### Surface plasmon resonance (SPR)

The affinity between DPP-IV and the selected peptides was evaluated by using a Biacore 8K high-throughput intermolecular interaction analysis system (GE Healthcare, Chicago, IL) with CM5 chips (Cytiva, Marlborough, MA) at 25 °C. The specific experimental processes were performed according to our previous study (*3*).

### Simulated digestive stability of peptides

The *in vitro* digestibility of the peptides was determined according to the method described by Ohanenye *et al*.(*55*) with some modifications. Briefly, 5.0 mg/mL peptides were mixed with 5000 U/mL porcine pepsase (pH 3.0). The mixture was incubated with continuous shaking in a water bath at 37 °C for 2 h. Then, the mixture was adjusted to pH 7.0, and a final concentration of 100 U/mL trypsase was added, followed by continuous shaking at 37 °C for another 2 h. After digestion, the enzyme was inactivated, and the mixture was incubated in a water bath (100 °C) for 20 min. Subsequently, the mixture was centrifuged at 6000 rpm for 20 min, after which the supernatants were collected for further analysis. The concentrations of peptides in the digestions and TPH were determined via AB SCIEX QTrap 6600+ (AB Sciex, America).

### Intestinal organoid model for evaluating DPP-IV inhibition

To evaluate DPP-IV inhibition, organoids were treated with 500 μg/mL peptides for 24 h, the mixtures were centrifuged at 500 × g for 5 min, and the supernatants were used to determine the DPP-IV inhibition rate and concentrations of active GLP-1 by using a DPP-IV-Glo^TM^ Protease Assay kit and GLP-1 ELISA Kit, respectively.

### Animals and treatments

All our animal experiments were approved by the Shenzhen Bay Laboratory (permit: AEXXH202201). C57BL/6J mice were purchased from Beijing Vital River Laboratory Animal Technology Co., Ltd. (Beijing, China). *ob/ob* mice (T001461) were purchased from GemPharmatech Laboratory (GemPharmatech, Nanjing, China). Mice were maintained under a strict 12 h light cycle and had unlimited access to water and food. All mice were randomly assigned to the experimental groups; the groups did not present differences in body weights before the treatments, and no mice were excluded from the analysis.

To evaluate the hypoglycaemic activity of TPH, all mice were fed a high-fat diet (HFD, Research Diets, cat# D12492). Eight-week-old C57BL/6J SPF mice were supplemented daily with PBS (HFD group), sitagliptin (12.5 mg/kg, Sit group), hemp seed protein hydrolysates by thermolysin (80 mg/kg, TPH group), and hemp seed protein (320 mg/kg, HSP group) for 8 weeks by oral gavage.

To test the *in vivo* effect of VAMP on host DPP-IV, 8-week-old C57BL/6J SPF mice were fed a HFD for 8 weeks, then treated daily with VAMP (50 mg/kg) or PBS (vehicle group) for 1 week.

To investigate the effects of VAMP on glucose metabolism, 8-week-old C57BL/6J SPF mice were treated daily with PBS (vehicle group) or VAMP (50 mg/kg, VAMP group) for 8 weeks by oral gavage.

To test the hypoglycaemic and DPP-IV inhibitory effects of VAMP in *ob/ob* mice, 6-week-old *ob/ob* mice were treated daily with PBS (vehicle group) or VAMP (50 mg/kg, VAMP group) for 4 weeks by oral gavage.

For the preliminary safety evaluation, C57BL/6J male SPF mice were divided into three groups. The treatment groups were given 20 mg/kg or 100 mg/kg VAMP 3 times per week by gavage. The vehicle groups were given an equivalent volume of PBS by oral gavage. The treatment continued for 4 weeks.

For FMT experiment, mice were treated with Abx for one week. Feces from mice treated with VAMP or PBS were collected. Feces (100 mg) were resuspended in sterile anaerobic PBS (1 mL) and then centrifuged at 200 × g for 3 min at 4 °C, the supernatant was collected and administered to mice after Abx treatment.

To test the effects of *A. muciniphila* on glucose metabolism in HFD-fed mice, mice were fed a HFD for 8 weeks, then treated with *A. muciniphila* or PBS 3 times per week for 4 weeks. For bacterial colonization, mice were given *A. muciniphila* by gavage at a dose of 2 × 10^8^ CFUs/0.2 mL suspended in sterile anaerobic PBS.

To investigate the effects of *A. muciniphila* in improving glucose metabolism during VAMP treatment, mice were fed a HFD for 8 weeks. Then mice were treated with PBS, 50 mg/kg VAMP, 50 mg/kg benzydamine hydrochloride, or 50 mg/kg VAMP combined with benzydamine hydrochloride 3 times per week for 4 weeks.

### Metabolic assays

Oral glucose tolerance tests (OGTTs) were conducted following a 6 hour fasting period in mice. Blood glucose levels were assessed using a glucometer on tail vein blood samples at 0, 15, 30, 60, and 90 minutes after oral glucose administration with a dosage of 1.5 g/kg body weight. For insulin tolerance tests (ITTs), mice were intraperitoneal injected with insulin (0.8 U/kg body weight) after 6 hours of fasting. For measurement of active GLP-1 (Millipore, cat# EGLP-35K), total GLP-1 (Elabscience, cat# E-EL-M0090c), and insulin (ABclonal, cat# RK02951), blood was collected before gavage and 15 min after glucose gavage; the blood was supplemented with 10 mM sitagliptin, and the plasma was stored at −80 °C until further analysis. To measure the indicators in intestinal tissue, a 0.5-cm segment of distal gut tissue was homogenized in RIPA buffer supplemented with 10 mM sitagliptin. The levels of active GLP-1, total GLP-1, and insulin were measured using the corresponding kits and normalized to the protein concentration (via a BCA protein assay).

### Plasma biochemical analysis

The levels of plasma total cholesterol (TC), triacylglycerol (TG), high-density lipoprotein cholesterol (HDL-c), low-density lipoprotein cholesterol (LDL-c), nonesterified fatty acid (NEFA), tumour necrosis factor α (TNF α), and interleukin-1β (IL-1β) were determined by commercial kit (Nanjing Jiancheng Bioengineering Institute, Jiangsu, China).

### Intestinal permeability analysis

Intestinal permeability was evaluated through the oral administration of fluorescein-isothiocyanate (FITC)-dextran in mice that were subjected to a 4-hour fasting period. Following the gavage of FITC-dextran at a dose of 200 mg/kg body weight, blood samples were collected from the tail vein after 90 minutes. The collected blood was centrifuged at 3000 × g for 10 minutes to obtain serum samples. Subsequently, 20 μL aliquots of serum were plated in 96-well plates, diluted with PBS to a final volume of 200 μL, and analyzed for fluorescence intensity at excitation and emission wavelengths of 485 nm and 520 nm, respectively.

### Gene expression analysis

Mouse tissues were cryopreserved in liquid nitrogen at −80 °C, followed by standard phenol‒chloroform extraction using TRIzol reagent to isolate total RNA. Subsequently, cDNA was synthesized from 2 μg of total RNA using a reverse transcription kit. The quantification of individual mRNA levels was determined by normalizing to β-actin mRNA.

Reverse transcription of total RNA was performed using the PrimeScript RT reagent kit (Takara). Quantitative real-time PCR was performed using TB Green Premix Ex Taq II (Takara) on a QuantStudio 7 Flex Real-Time PCR system (Thermo Fisher Scientific, Waltham, MA, USA). The primers were listed in the Key Resources Table. β-actin was used as an endogenous control. The relative mRNA expression levels were calculated using the 2^−△△Ct^ quantification method.

Mouse tissues were frozen and stored at −80 °C, then RNA was extracted from the tissue using RNA isolation Kit. cDNA was synthesized using QuantiTect Reverse Transcription kits. Quantitative real-time PCR was performed using TB Green Premix Ex Taq II (Takara) on a QuantStudio 7 Flex Real-Time PCR system (Thermo Fisher Scientific, Waltham, MA, USA). β-actin was used as an endogenous control. The relative mRNA expression levels were calculated using the 2^−△△Ct^ quantification method. The sequences of primers used for RT‒qPCR are shown in **Table S3**.

### Histological analysis

For histological analysis of subcutaneous white adipose tissue, subsections were partially embedded in 10% neutral buffered formalin solution (Sigma, USA). Paraffin-embedded adipose tissue sections were stained with haematoxylin and eosin (H&E) for morphological examination. Changes in histological fat sections were observed under a light microscope, and the cell area was calculated with ImageJ software (ImageJ Software Inc., USA).

For histological analysis of colonic tissue, colon segments were promptly immersed in Carnoy’s fixative for 24 hours. Subsequent, the colon specimens underwent standard dehydration procedures before being embedded in paraffin, and thin sections (5 μm) were cut and deposited on glass slides. The paraffin sections were stained with Alcian blue/periodic acid-Schiff. The thickness of the colonic sections was then measured by ImageJ, and 5 different measurements were made perpendicular to the inner mucus layer per image. Only regions in which the mucus layer was sandwiched between the epithelium on one side and luminal contents on the other were used; care was taken to measure regions that represented the average thickness in each blinded image.

### Gut microbiota analysis

Total DNA was extracted from colonic contents using a Tiangen stool DNA extraction kit. The V3-V4 region of the 16S rRNA gene was amplified by PCR from the extracted and purified genomic DNA using 515 forward and 806 reverse primer pairs. PCR amplification was performed on a PCR System (Bio-Rad, Hercules, CA, USA), and the PCR amplification products were separately extracted from a 2% agarose gel and further purified using a Tiangen agarose gel DNA purification kit. The purified amplicons were quantified using a Qubit Fluorometer (Thermo Fisher Scientific, Waltham, MA, USA), pooled in equimolar, and sequenced on an Illumina Miseq platform (Illumina, San Diego, CA, USA).

Raw sequencing data were processed with QIIME2. In brief, the forward and reverse reads each were truncated at 200 bases. Taxonomy was assigned using the Greengenes reference (version 13.8) database. Analyses of alpha diversity (one-way ANOVA followed by Tukey’s post hoc test), beta diversity (weighted UniFrac distance followed by ANOSIM test), and bacterial taxonomic distributions were performed using MicrobiomeAnalyst.

### Stability of the VAMP

The faeces-derived bacterial community or *A. muciniphila* were incubated in BHI medium supplemented with 100 μM VAMP for 48 h. The levels of VAMP at different time points were quantified by targeted metabolomics.

### Statistical analysis

All the statistical data were analysed using SPSS version 26.0. All experimental data are reported as the mean ± SEM. Student’s t test or one-way ANOVA for multiple comparisons followed by Tukey’s test was used to determine the significance of differences. *p* < 0.05 was considered significant.

## Acknowledgements

Financial support was received from the Shenzhen Science and Technology Plan Project (Grant No. JCYJ20230807111614030), Youth Science Foundation Project (Grant No. 32101936), Shenzhen Science and Technology Innovation Commission (Grant No. KCXFZ20201221173207022), and China Postdoctoral Science Foundation, No.15 Special funds (In-Station) (Grant No. 2022T150366), Hechi Research and Development Program (Grant No. HekeAA230807) are gratefully acknowledged.

## Author contributors

Conceptualization, H.C., Q.N. and X.X.; methodology, H.C., Q.N., W.L., Q.Z., Y.W., J.L.; software, H.C.; investigation, H.C.; resources, H.C. and X.X.; writing – original draft, H.C. Q.N., and X.X.; writing – review & editing, H.C., Q.N., W.L., W.H., B.X., Y.W., J.L., C.Z., X.Z., X.X.; visualization, H.C.; supervision, H.C., Q.N., Y.W. and X.X; project administration, X. X; funding acquisition, H.C. and X.X.

## Data and materials availability

All data are available in the main text or the supplementary materials.

## Supplementary Materials

**Figure S1.**
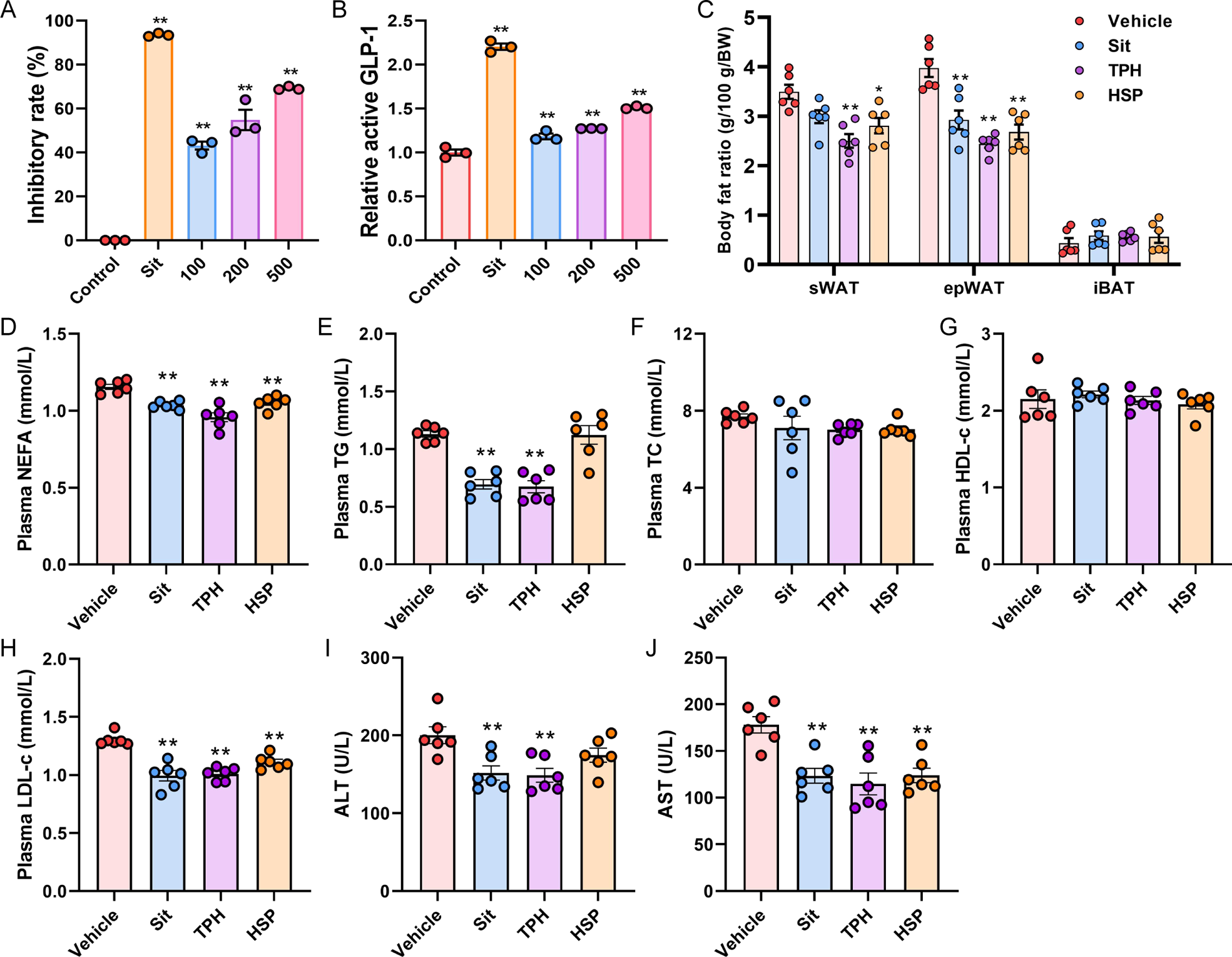
Treatment with TPH improved host metabolism. (A) The inhibitory effect of different concentrations of TPH on DPP-IV activity was measured. (B) The relative concentrations of active GLP-1 after TPH treatment in intestinal organoids were determined. (C to J) HFD-fed mice were treated with PBS (vehicle group), sitagliptin (Sit group), HSP hydrolysates from thermolysin (TPH group), or HSP (HSP group) for 8 weeks by oral gavage. n = 6 mice/group. (C) The weights of sWAT, epWAT, and iBAT were determined. (D-J) The levels of plasma NEFAs, TG, TC, HDL-c, LDL-c, ALT, and AST were determined. All the data are presented as the mean ± SEM. \**P* < 0.05 and \*\**P* < 0.01 versus the vehicle group.

**Figure S2.**
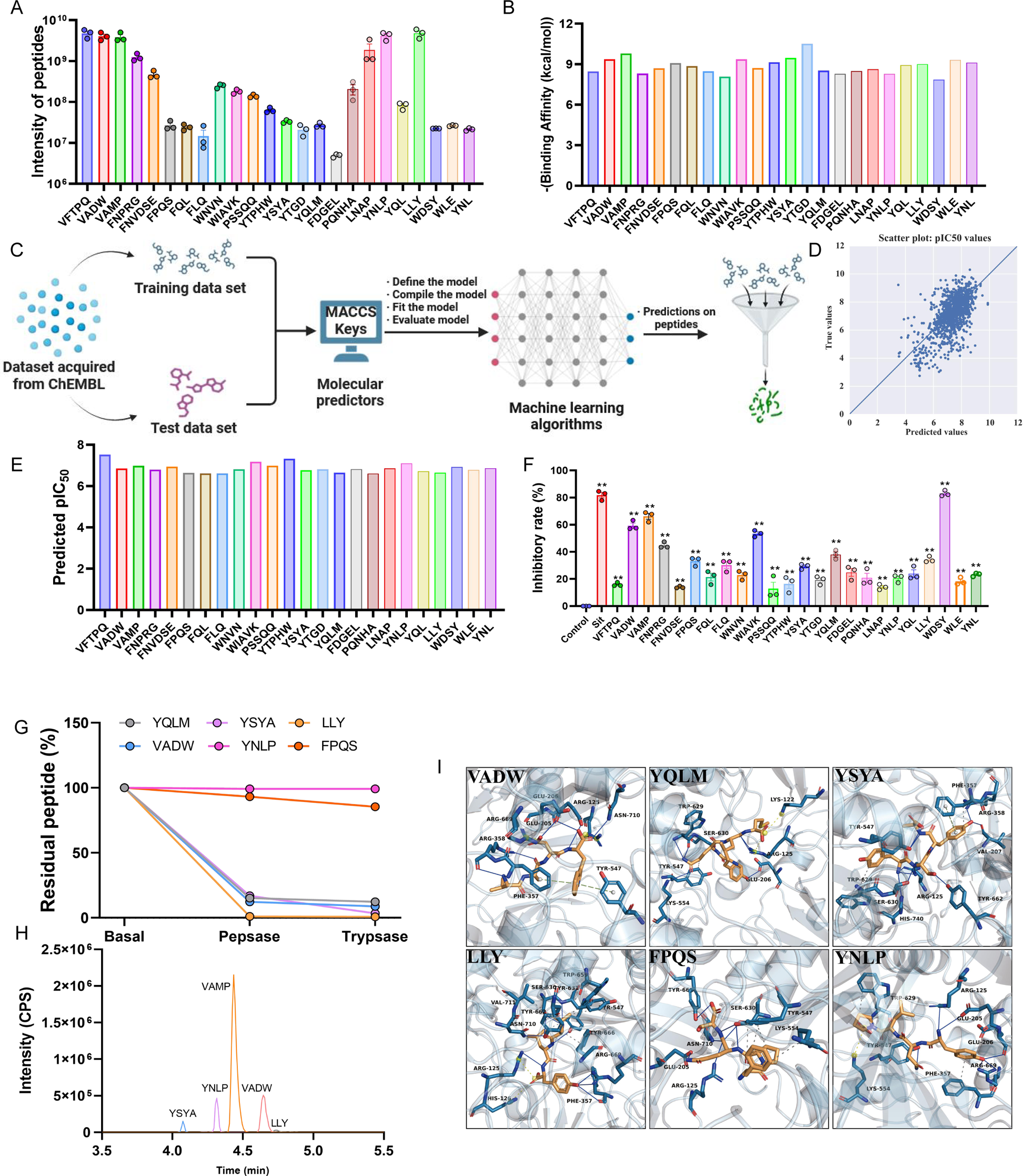
VAMP is an effective peptide inhibitor of DPP-IV. (A) The relative intensity of the 24 selected peptides in the TPH sample is shown. (B) PyRx-Dock-derived docking binding affinity for DPP-IV (receptor) and the 24 selected peptides. (C) Strategy for machine learning. (D) Scatter plot for visualization of the correlation between the predicted and true pIC_50_ values in the test set. (E) Predicted pIC_50_ values of the 24 selected peptides. (F) DPP-IV inhibition by the 24 selected peptides. (G) The residual VADW, YQLM, YSYA, LLY, FPQS, and YNLP after pepsase and trypsase digestion are shown. (H) Representative extracted ion chromatograms of YSYA, YNLP, VAMP, VADW, and LLY in the TPH sample are shown. (I) Active-site superimposition between peptide (VADW, YQLM, YSYA, LLY, FPQS, and YNLP)-bound DPP-IV is shown with key residues labelled with three-letter codes. Peptides (claybank) and residues (navy blue) are shown as sticks, and DPP-IV is shown as a cartoon. Hydrogen bonds are coloured blue, hydrophobic interactions are coloured grey, salt bridges are coloured croci, and π-stacking interactions are coloured light green.

**Figure S3.**
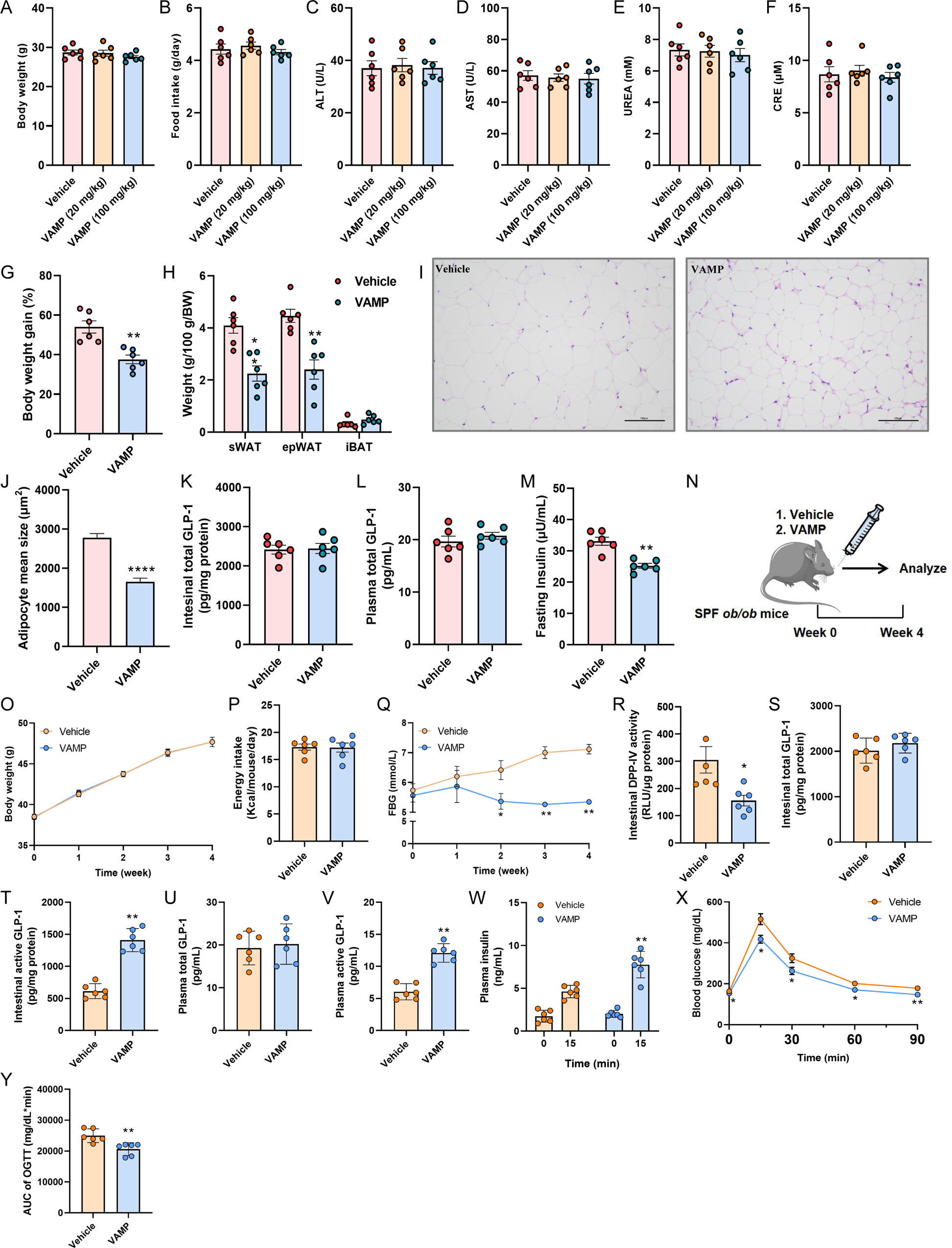
Safety and hypoglycaemic effects of VAMP. (A to F) SPF mice were divided into three groups and treated with 20 mg/kg VAMP, 100 mg/kg VAMP or vehicle for 4 weeks; n = 6 mice/group. (A) The body weights, (B) food intake, (C) ALT levels, (D) AST levels, (E) UREA levels, and (F) CRE levels were determined. (G to M) HFD-fed mice were treated with PBS (vehicle group) or VAMP (50 mg/kg, VAMP group) for 8 weeks by oral gavage. n = 6 mice/group. (G) The body weight gain and (H) weights of sWAT, epWAT, and iBAT are shown. (I) Representative H&E images of sWAT deposits (3 mice per group) are shown. Scale bars, 100 μm. (J) The mean adipocyte size (3 mice per group), (K) intestinal total GLP-1 levels, (L) plasma total GLP-1 levels, and (M) fasting insulin levels were determined. (N) The experimental scheme for (O to Y) is shown, n = 6 mice/group. The *ob*/*ob* mice were divided into two groups and treated with PBS (vehicle group) or VAMP (50 mg/kg, VAMP group) for 4 weeks by oral gavage. (O) Body weight changes, (P) energy intake, (Q) changes in FBG levels, (R) DPP-IV activity in intestinal tissue, (S) intestinal total GLP-1 levels, (T) intestinal active GLP-1 levels, (U) plasma total GLP-1 levels, (V) plasma active GLP-1 levels, (W) glucose-stimulated insulin levels, (X) OGTT curve, and (Y) AUC of the OGTT were evaluated. All the data are presented as the mean ± SEM. \**P* < 0.05 and \*\**P* < 0.01 versus the vehicle group.

**Figure S4.**
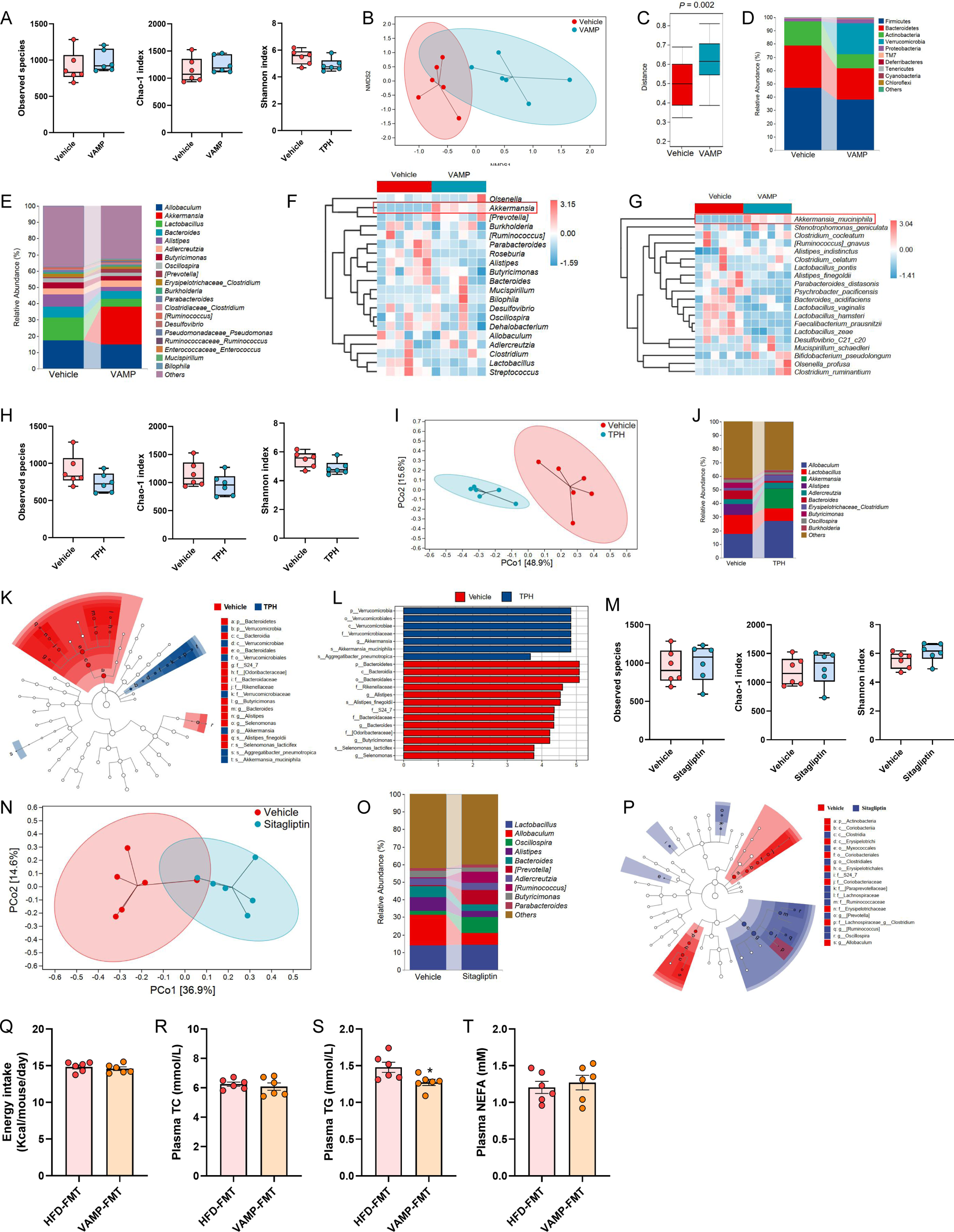
VAMP treatment increased the abundance of *A. muciniphila*. (A-G, related to Figure 3L) HFD-fed mice were treated with PBS (vehicle group) or VAMP (50 mg/kg, VAMP group) for 8 weeks by oral gavage. (A) The α diversity of the gut microbiota was compared between the vehicle and VAMP groups, as indicated by the observed species, Chao1, and Shannon indices. (B) Nonmetric multidimensional scaling (NMDS) analysis was performed. (C) The Bray ‒ Curtis distance. (D) Phylum-level compositions of the gut microbiota in the Vehiclevehicle and VAMP groups. (E and F) The genus-level compositions and heatmap of gut microbiota between the vehicle and VAMP groups. (G) A heatmap of the gut microbiota at the species level in the vehicle and VAMP groups is shown. (H-L, related to Figure 1) HFD-fed mice were treated with PBS (vehicle group) or TPH (TPH group) for 8 weeks by oral gavage. (H) The α diversity of the gut microbiota was compared between the vehicle and TPH groups, as indicated by the observed species, Chao1, and Shannon indices. (I) PCoA was performed using the Bray‒Curtis distance. (J) The genus-level differences in the composition of the gut microbiota between the vehicle and TPH groups were evaluated. (K-L) Taxonomic cladograms were generated by LEfSe analysis. The blue colour indicates enriched taxa in the TPH group, and the red colour indicates enriched taxa in the vehicle group. The size of each circle is proportional to the taxon’s abundance. (M-P, related to Figure 1) HFD-fed mice were treated with PBS (vehicle group) or sitagliptin (sitagliptin group) for 8 weeks by oral gavage. (M) The α diversity of the gut microbiota was compared between the vehicle and sitagliptin groups, as indicated by the observed species, Chao1, and Shannon indices. (N) PCoA was performed using the Bray ‒ Curtis distance. (O) The genus-level composition of the gut microbiota was compared between the vehicle and sitagliptin groups. (P) Taxonomic cladograms were generated by LEfSe analysis. The blue colour indicates enriched taxa in the Sitagliptin group, and the red colour indicates enriched taxa in the Vehicle group. The size of each circle is proportional to the taxon’s abundance. (Q) The energy intake of mice in the HFD-FMT and VAMP-FMT groups was measured. (R) Plasma TC content, (S) TG content, and (T) NEFA content were determined. All the data are presented as the mean ± SEM. \**P* < 0.05 and \*\**P* < 0.01 versus the HFD-FMT group.

**Figure S5.**
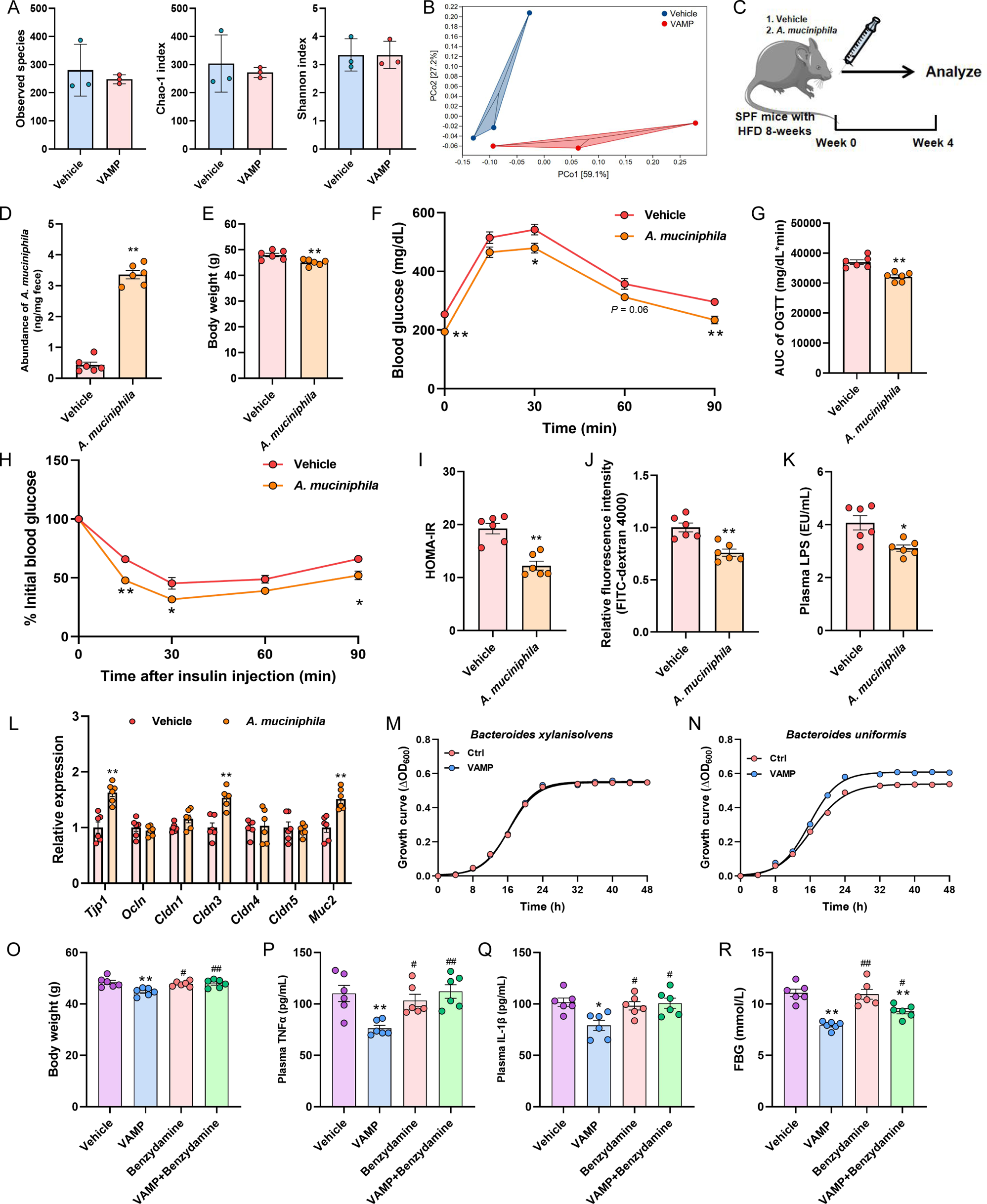
*A. muciniphila* is an important factor that mediates the metabolic protective effect of VAMP. (A-B) Mouse faeces-derived *ex vivo* communities were cultured in BHI media supplemented with VAMP (200 μM) or PBS and then grown anaerobically for 48 h. (A) The α diversity of the gut microbiota was compared between the vehicle and VAMP groups, as indicated by the observed species, Chao1, and Shannon indices. (B) PCoA was performed using the Bray‒ Curtis distance. (C) The experimental scheme for D to L is shown, n = 6 mice/group. Mice were fed a HFD for 8 weeks and then treated with PBS (vehicle group) or *A. muciniphila* (*A. muciniphila* group) two times per week for 4 weeks. (D) *A. muciniphila* abundance in faeces was assessed by qPCR. (E) The body weight, (F) OGTT curve, (G) AUC of the OGTT, (H) ITT curve, and (I) HOMA-IR index in different groups were evaluated. (J) Intestinal permeability was measured by plasma fluorescence intensity after gavage with FITC-dextran 4000. (K) The plasma LPS level was evaluated. (L) The relative expression of *Tjp1*, *Ocln*, *Cldn1*, *Cldn3*, *Cldn4*, *Cldn5* and *Muc2* mRNAs in colonic tissue was determined. (M-N) The growth curves of *Bacteroides xylanisolvens* and *Bacteroides uniformis* cultured in BHI media supplemented with VAMP (50 μM) or PBS control are shown; n = 3 biological replicates. (O-R) Mice were fed a HFD for 8 weeks and then treated with PBS (vehicle group), VAMP (50 mg/kg, VAMP group), benzydamine (50 mg/kg, benzydamine group), or VAMP plus benzydamine (50 mg/kg, VAMP+benzydamine group) three times per week for 4 weeks. (O) The body weight, (P) plasma concentrations of TNFα and (Q) IL-1β, and (R) FBG level were determined. All the data are presented as the mean ± SEM. In D-L, \**P* < 0.05 and \*\**P* < 0.01 versus the vehicle group. In O-R, \**P* < 0.05 and \*\**P* < 0.01 versus the vehicle group; #*P* < 0.05 and ##*P* < 0.01 versus the VAMP group.

**Table S1.** Amino acid contents in HSPs.

| Amino acids | Contents, $\mu\text{g/g}$, n=3 |
| --- | --- |
| L-arginine | 2496.28±10.73 |
| L-aspartic acid | 2323.48±4.47 |
| L-cystine | 34.85±1.16 |
| L-glutamic acid | 3404.301±14.93 |
| Glycine | 790.68±0.91 |
| L-histidine | 552.82±0.96 |
| L-leucine | 1124.01±1.35 |
| L-Lysine | 548.225±0.73 |
| L-methionine | 212.89±4.46 |
| L-phenylalanine | 968.53±0.06 |
| L-proline | 568.79±1.48 |
| L-serine | 954.88±3.58 |
| L-threonine | 481.98±0.93 |
| L-tyrosine | 723.08±1.29 |
| L-valine | 1272.86±8.70 |
| L-alanine | 797.54±1.21 |
| Hydroxyproline | 3.02±0.01 |
| Sarcosine | 142.16±1.84 |
| Homoproline | 164.69±1.84 |

**Table S2.** Conditions for protein hydrolysis.

| Enzyme | temperature<br>(°C) | pH | Enzymatic activity (U/g) |
| --- | --- | --- | --- |
| Trypsase | 45 | 8 | 10000 |
| Pepsase | 45 | 2.5 | 10000 |
| Papain | 55 | 7 | 10000 |
| Chymotrypsin | 55 | 7 | 10000 |
| Alcalase | 55 | 10.5 | 10000 |
| Flavourzyme | 55 | 7 | 10000 |
| Neutral protease | 55 | 7 | 10000 |
| Proteinase K | 55 | 7 | 10000 |
| Thermolysin | 55 | 7 | 8000 |
| Trypsase/Pepsase | 40 | 2.5/8.0 | 10000 |
| Trypsase/Chymotrypsin | 55 | 7.0/8.0 | 10000 |

**Table S3.**
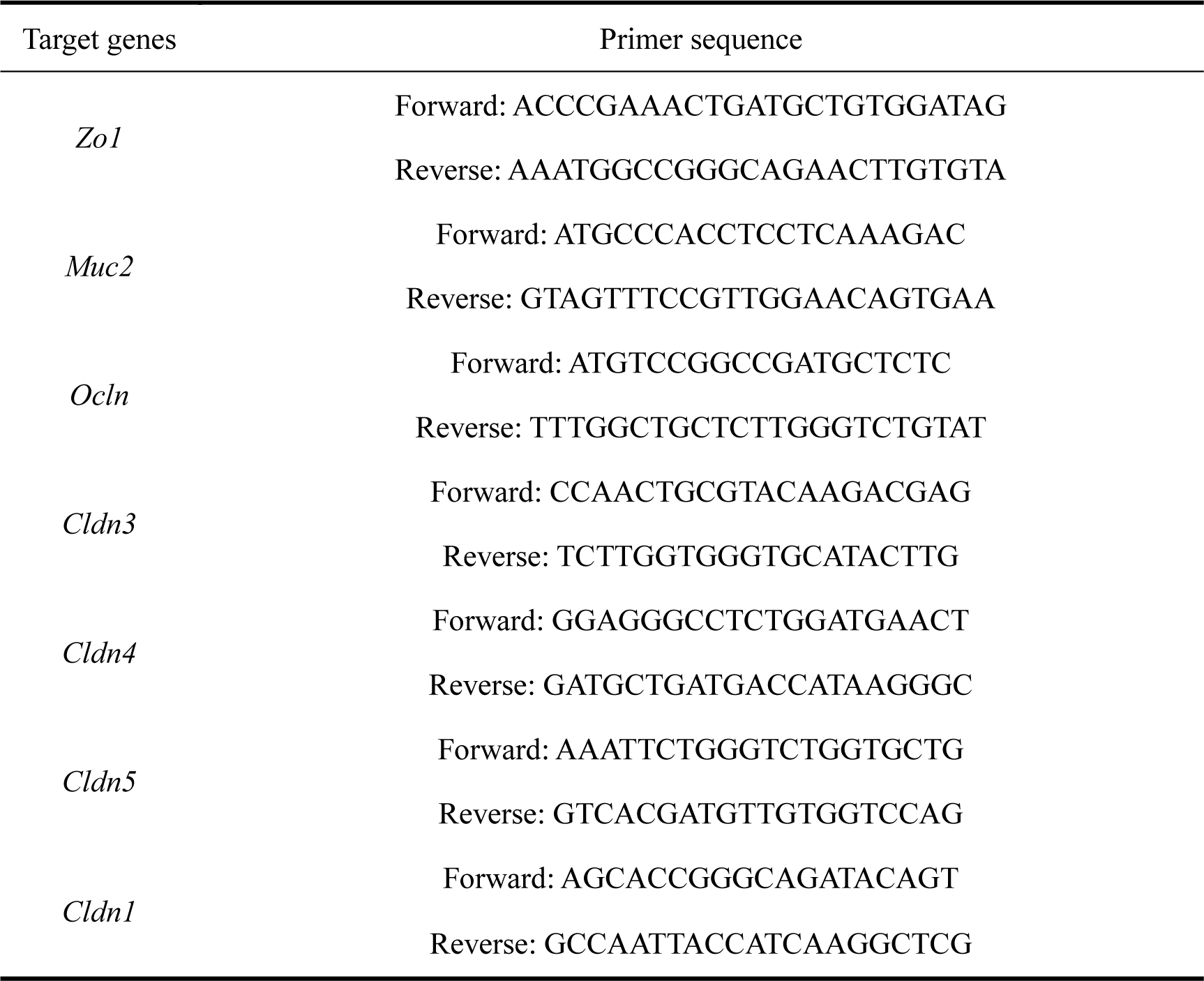
Primer sequences for RT‒qPCR.

